# Intra-Tumoral Treg Ablation Unleashes NK Cell-Mediated Control of CD8 T Cell-Resistant Tumors

**DOI:** 10.1101/2025.06.26.661417

**Authors:** Chenyu Zhang, Charles Chien, Eglė Jurgaitytė, Koharu Sakiyama, Alissa Bockman, Yeara Jo, Seungwon Lee, Stephanie Silveria, Elizabeth Andrews, Abigail Mende, Lily Zhang, K. Christopher Garcia, Allon Wagner, Michel DuPage, David Raulet

## Abstract

Cancer cells frequently lose MHC I to evade CD8^+^ T cell recognition. While Natural Killer (NK) cells are poised to target MHC I-deficient cancer cells, MHC I loss alone is often insufficient to unleash fully effective NK cell responses. Here we show that selective intra-tumoral (IT) ablation of regulatory T (Treg) cells elicited potent antitumor NK cell responses that controlled MHC I-deficient and even MHC I^+^ cancers. Tregs controlled the activation, maturation, and anti-tumor cytotoxic activity of NK cells within the tumor microenvironment. Mechanistically, IT Tregs prevented the cDC2-dependent induction of IL-2 production by CD4^+^ Tconv cells that was necessary for NK cell activation. Systemically administered antibodies that selectively depleted IT Tregs similarly empowered NK-dependent tumor control. These findings expand the breadth of Treg-mediated cancer immunosuppression to encompass antitumor NK cells and suggest that targeting Tregs in tumors can control CD8^+^ T cell-resistant cancers.

**One Sentence Summary:** IT Treg ablation drives NK cell tumor control via CD4^+^ Tconv-derived IL-2, eliminating MHC I+/MHC I- cancers without systemic toxicity.

## Introduction

Downregulation or loss of MHC I is a frequent mechanism of tumor immune in human tumors^1^. MHC I loss is common in multiple tumor types and associated with mutations occurring in the genes encoding MHC proteins themselves, as well as many genes associated with antigen processing and presentation. By reducing or eliminating MHC I expression, tumor cells become resistant to being killed by CD8^+^ T cells, which recognize tumor antigens presented by MHC I molecules. Thus, tumor cells undergo strong selective pressure to lose MHC I antigen presentation to evade CD8-mediated immune control, which is especially apparent in the setting of checkpoint blockade immunotherapies that strengthen CD8^+^ T cell responses^2–4^.

Natural Killer (NK) cells are cytolytic lymphocytes that recognize transformed and virus-infected cells using germline-encoded receptors to recognize stimulatory ligands upregulated on unhealthy cells. Activated NK cells control cancer by directly killing tumor cells via perforin/granzyme secretion or death receptor engagement, or kill indirectly by producing cytokines (e.g. IFN-γ, TNF-α) and chemokines that enhance immune infiltration and activation at the site of the tumor^5,6^. Loss of MHC I enhances tumor cell killing by NK cells by preventing engagement of MHC I-specific inhibitory receptors on NK cells (“missing self” recognition). Nevertheless, loss of MHC I alone is frequently insufficient to elicit tumor-controlling NK cell responses^7^, since MHC I- deficient tumors typically progress in the absence of therapeutic interventions. However, recent studies demonstrate that agents that mobilize and activate NK cells can effectively treat multiple tumor types, including established solid tumors^7–9^. We hypothesize that without intervention, NK cell anti-tumor responses are held in check by immunosuppressive mechanisms or the absence of sufficient activation signals for NK cells.

In normal tissues, Tregs maintain tissue homeostasis by preventing inappropriate or excessive immune responses while also directly promoting tissue repair. How Tregs are recruited to tumors and how they protect tumor cells from elimination by the immune system are active areas of investigation^10^. Early Treg-based cancer therapy studies using anti-CD25 and anti-CTLA-4 antibodies to deplete Tregs demonstrated rejection of transplanted tumors^8,9,11–15^. However, understanding the impact of each antibody is difficult due to their effects on other immune cells, especially tumor-controlling effector T cells, as well as their impact systemically. We previously generated genetically engineered mouse models of cancer, along with approaches to disrupt Treg functions locally within tumors, to study the impact of removing immunosuppression on anti-tumor immunity, primarily by CD8^+^ T cells^16,17^. In normal non-tumor bearing mice, the depletion of Tregs using the *Foxp3^DTR-GFP^* system results in massive expansion and activation of many immune cell types including CD4^+^ and CD8^+^ T cells, B cells, granulocytes, dendritic cells (DCs), and NK cells^18–22^. Tregs execute their suppressive activity by several different mechanisms^10,18,19,23–27^. It is known that due to their high expression of IL-2Rα (CD25), Tregs reduce the availability of IL-2 to act on other T cells, serving as an IL-2 sink, while also suppressing the production of IL-2 by other T cells^27–30^. Inhibition of NK cell proliferation and function by Tregs has been described in homeostatic or autoimmune conditions using systemic Treg depletion. However, the impact of Treg depletion on antitumor NK cell responses, especially in the setting of local Treg depletion within the tumor environment, has not been investigated in depth^24,29–32^. Previous studies have highlighted Treg suppressive mechanisms by the secretion of inhibitory cytokines (TGF-β, IL-10, IL-35, IL-37), their expression of CTLA-4 and PD-L1, and their capacity to block key interactions of NK cells with DCs and macrophages. Contact-dependent inhibition of NK cells by Tregs can occur by cell surface-bound TGF-β on Tregs^24^.

Even less is known about the role of Tregs in suppressing antitumor NK cell responses locally within the tumor microenvironment. Here, we show that selective IT depletion of Tregs unleashes a powerful NK cell response locally within tumors that controls the growth of MHC I-deficient tumors without inciting autoimmunity. Local NK cell activation, proliferation, and cytolytic function required CD4^+^ T cells that produced IL-2. IL-2 was both necessary for tumor control, and sufficient to restore NK cell antitumor activity if CD4^+^ Tconv cells were depleted in combination with Tregs. In addition, conventional type 2 dendritic cells (cDC2s) were critical for NK-mediated tumor control, likely because they were required to prime CD4^+^ Tconv cells to produce IL-2 in tumors after IT Treg ablation. Finally, therapeutically translatable approaches to deplete IT Tregs with antibodies led to tumor control in a manner that recapitulated our genetic *Foxp3^DTR^* experimental model. Taken together, our findings show that IT Treg ablation activates a NK-CD4-DC axis that promotes NK cell-mediated control of tumors, even against tumors that are resistant to CD8^+^ T cells.

## RESULTS

### Intra-tumoral Treg ablation controls MHC I-deficient tumors without autoimmunity

While Tregs have been shown to restrain T cell responses to MHC I+ tumors, less is known about local Treg-mediated suppression of anti-tumor NK cell responses. We engineered three different C57Bl/6 (B6) MHC I-deficient tumor models generated by CRISPR-Cas9-mediated disruption of the beta-2-microglobulin (*B2m)* gene: the MC38- *B2m^-/-^* (colon carcinoma), B16F10-*B2m^-/-^* (melanoma) and RMA-*B2m^-/-^* (lymphoma) cell lines^33,34^ (Ext. Data Fig. 1a). We depleted Tregs using *Foxp3^DTR-GFP^* mice, wherein diphtheria toxin (DT) injection leads to Treg apoptosis^18,33,34^. First, mice bearing subcutaneous tumors of ∼50 mm^3^ (five days after inoculation) began receiving DT systemically every other day by intraperitoneal (IP) injection (Fig. 1A). IP DT treatments resulted in Treg depletion in all organs assessed and the significant control of all three *B2m^-/-^* tumor types (Fig. 1B-C, Ext. Data Fig 1c-d). These results highlight that Treg ablation can lead to tumor control even when tumors lack MHC I molecules. However, systemic Treg depletion led to autoimmunity, as evidenced by weight loss (Fig. 1D, Ext. Data Fig. 1b), enlarged lymph nodes (Ext. Data Fig. 1e), increased spleen weight (Fig. 1E), and morbidity by day 15 that required euthanasia.

**Figure 1.**
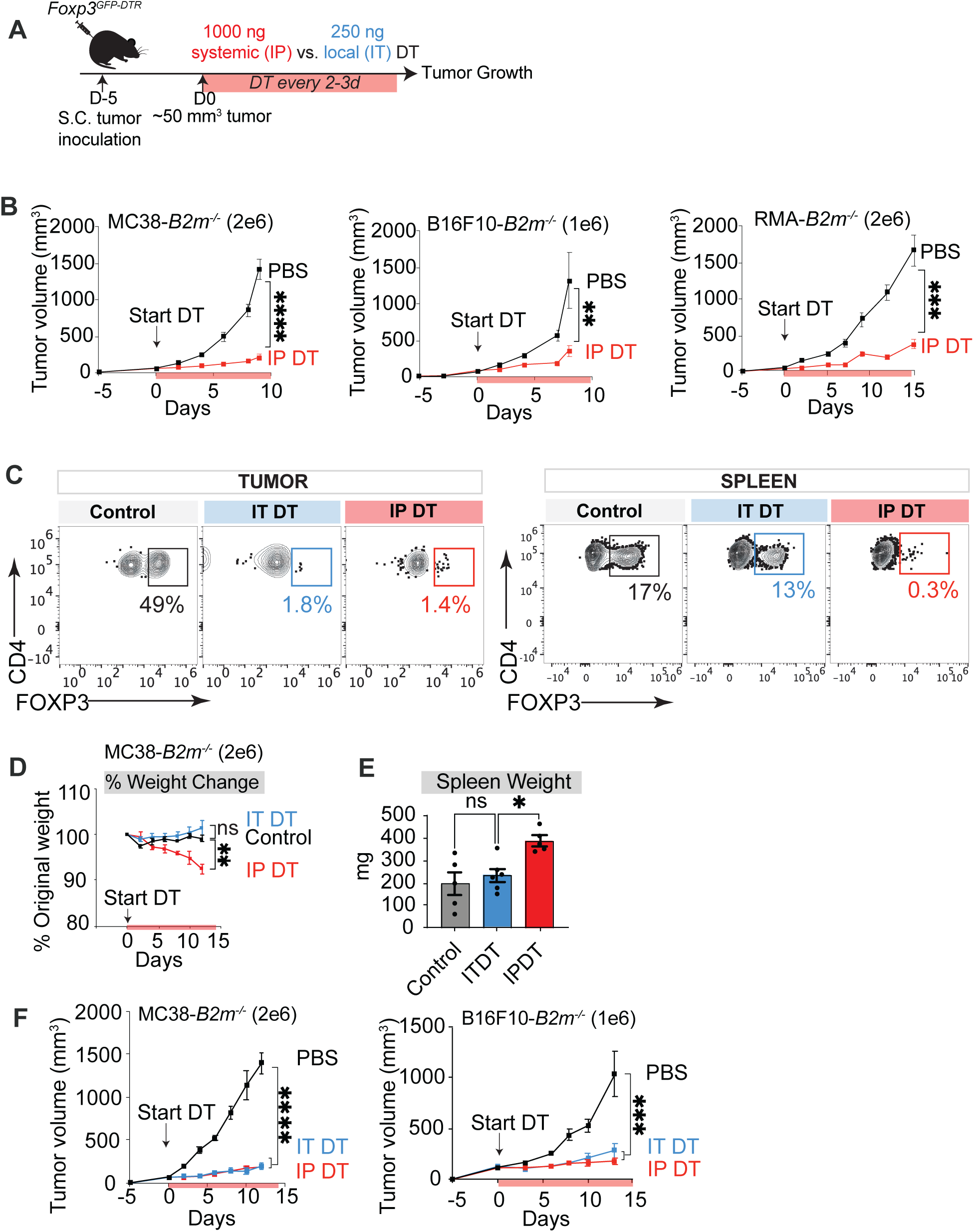
Intra-tumoral Treg depletion leads to potent tumor control without morbidity (A) Schematic of tumor inoculation and treatment regimen in *Foxp3^DTR-GFP^* mice depleted of Tregs either systemically by IP injections of 1000 ng DT or locally (intra-tumorally, IT) with 250 ng of DT. DT inoculations began when tumors reached ∼50 mm^3^, typically 5 days after tumor inoculations. (B) Tumor growth curves in *Foxp3^DTR-GFP^* mice systemically depleted of Tregs (IP DT). Data are representative of 3-10 experiments performed for each tumor type. (C) Representative flow plots showing DT-induced depletion of Tregs in tumors but not spleens of IT DT-treated MC38-*B2m^-/-^* mice on day 12 after DT treatment (blue boxes). Tregs were depleted in both sites in mice administered DT IP (red boxes). Gated CD4+CD3+ cells are shown. Tregs were identified as intracellular FoxP3+ cells. (D) Body weight changes in *Foxp3^DTR-GFP^* mice bearing MC38-*B2m^-/-^* tumors that were depleted of Tregs systemically (IP DT) or intratumorally (IT DT) compared to no depletion (control). (E) Spleen weight of *Foxp3^DTR-GFP^* mice bearing MC38-*B2m^-/-^* tumors at D12 post DT initiation. (F) Tumor growth curves in *Foxp3^DTR-GFP^* mice that were depleted of Tregs intra-tumorally with IT DT injections. MC38-*B2m^-/-^* and B16F10-*B2m^-/-^* tumors were tested. Data are representative of 2 experiments performed for each tumor type. Data represent means ± SEM; *p < 0.05, **p < 0.01, ***p < 0.001, and ****p < 0.0001. B, D, F: RM two-way ANOVA followed by Geiser-Greenhouse correction. (n=5-8 mice/group). E: Ordinary one-way ANOVA with Dunnett’s multiple comparison test.

To test whether localized Treg ablation within tumors could also mediate tumor control, but without autoimmune toxicity, we tested a low dose of DT injected intra- tumorally (IT) in an attempt to generate localized intra-tumoral-specific Treg depletion (Fig. 1A). Compared to IP DT, IT DT treatment resulted in a similar near complete depletion of Tregs within tumors, but retained abundant Tregs in the spleen and lymph nodes, indicative of significant tumor-specific Treg depletion (Fig. 1c, Ext. Data Fig. 1e, f). In both tumor models tested (B16F10-*B2m^-/-^* and MC38-*B2m^-/-^*), low-dose IT DT injections resulted in control of tumor growth that was equivalent to systemic Treg depletions by IP DT (Fig. 1F, Ext. Data Fig. 1d). However, mice treated with IT DT did not lose weight (Fig. 1D), did not exhibit enlargement of non-tumor-draining lymph nodes (ndLN) or spleens (Fig. 1E, Ext. Data Fig. 1e), and survived long-term. Furthermore, while the percentages of CD4^+^ Tconv cells and CD8^+^ T cells expressing the activation marker CD25 increased in the spleens and ndLN of IP DT-injected mice, IT DT-injected mice showed no significant increase in T cell activation in ndLN, despite increases in the tumor-draining lymph nodes (tdLN) (Ext. Data Fig. 1f). CD25^+^ CD4^+^ Tconv cells were increased in the tdLN of both types of mice, though more so in the IP-injected mice (Ext. Data Fig. 1f). Therefore, locally delivered IT DT elicited potent tumor control without resulting in systemic T cell activation and the onset of autoimmunity that accompanies systemic Treg ablation.

### NK cells and CD4^+^ Tconv cells control MHC I-deficient tumors upon intra-tumoral Treg ablation

To identify the effector cell types responsible for tumor rejection after IT Treg depletion, different effector cell types were depleted *in vivo* with antibody treatments starting one day before Treg depletion by IP or IT DT treatment in mice with 50 mm^3^ established tumors (Fig. 2A). With systemic (IP DT) (Fig. 2B-D) or intratumoral (IT-DT) (Fig. 2E-G, Ext. Data Fig 2a-c) Treg depletion, tumor control was diminished when all CD4^+^ Tconv cells or NK cells were depleted in all three tumor models tested. Importantly, however, CD8^+^ T cell depletion never reduced tumor control in any of the MHC I- deficient tumor models (Fig. 2B-D, Ext. Data Fig. 2d). Thus, local intra-tumoral depletion of Tregs was sufficient to mediate significant tumor control of MHC I-deficient tumors independently of CD8^+^ T cells. Notably, the numbers of NK or CD4^+^ Tconv cells in the tumors were not significantly increased after Treg depletion, suggesting a greater effect of Tregs on the functions of these cells within tumors as opposed to their recruitment or proliferation (Fig. 2H, Ext. Data Fig 2e).

**Figure 2.**
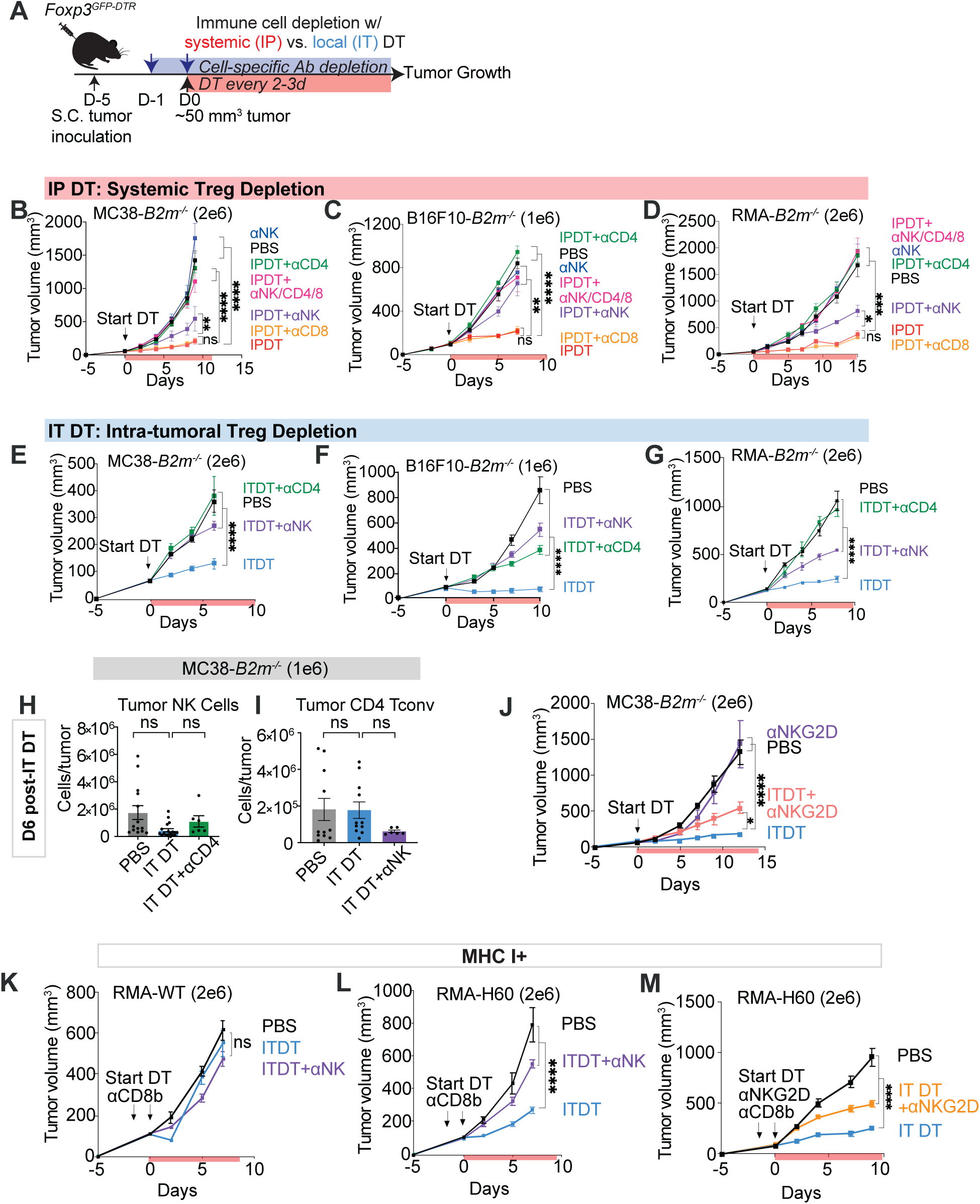
NK cells and CD4 T cells, but not CD8 T cells, are required for the control of tumors after Treg depletion. (A) Schematic of tumor inoculation and treatment regimen in *Foxp3^DTR-GFP^* mice depleted of Tregs systemically (IP) or locally (IT). DT was administered every 48-72 hours, while cell depleting antibodies were administered on D1, D0 and every six days thereafter. (B-D) Growth curves of various *B2m^-/-^* tumors in mice that were systemically depleted of Tregs alone (red) or in combination with depletion of all CD4 T cells (green), NK cells (blue), CD8 T cells (orange) or all three cell types (NK/CD4/CD8 cells) (pink). NK cell depletion alone without Treg depletion (purple) was included as an additional control. Data are representative of 2-3 experiments for each tumor type. (E-G) Growth curves of MC38-*B2m^-/-^,* B16F10-*B2m^-/-^* and RMA-*B2m^-/-^* tumors in mice that were intra-tumorally depleted of Tregs alone (blue) or in combination with depletion of all CD4 T cells (green) or NK cells (purple). Data are representative of 8 (MC38- *B2m^-/-^*), 3 (B16F10-*B2m^-/-^*) or 1 (RMA-*B2m^-/-^*) experiments. (H-I) Bar graphs showing percentages and counts of tumor NK cells and CD4 T cells six days after intra-tumoral Treg depletion (blue) or either Treg plus Tconv depletion (anti- CD4, green) or Treg plus NK depletion (anti-NK1.1, purple) in mice with MC38-*B2m^- /-^* tumors. Data are representative of 8 experiments. (J) Growth curves of MC38-*B2m^-/-^* tumors in mice depleted of Tregs with IT DT, without or with injections of blocking NKG2D antibody (clone MI-6) starting on D5. One experiment performed. (K-L) Growth curves of RMA-WT and RMA-H60 tumors in mice that were depleted of CD8b+ T cells before IT-Treg depletion without or with NK depletions with anti- NK1.1 antibody (clone PK136). One experiment performed. (M) Growth curves of RMA-H60a tumors in mice that were depleted of CD8b+ T cells before IT-Treg depletion without or with injections of blocking NKG2D antibody (clone MI-6). One experiment performed. Data represent means ± SEM; *p < 0.05, **p < 0.01, ***p < 0.001, and ****p < 0.0001. B, C, D, E, F, G, J-LM RM two-way ANOVA followed by Geiser-Greenhouse correction. (n=5-8 mice/group). H, I: Ordinary one-way ANOVA with Dunnett’s multiple comparisons test, representative of at least 2 independent experiments).

### IT Treg ablation elicits NK-dependent control of MHC I+ tumors expressing NKG2D ligands

Other than killing tumors cells via the loss of MHC I, i.e. “missing-self” recognition, NK cells also utilize activating natural cytotoxicity receptors (NCRs, e.g. NKG2D), which detect and bind to activating ligands often induced on cancer cells due to varying types of cellular stress^35–40^. MC38 tumors are known to naturally express NKG2D ligands^41^. Therefore, we used anti-NKG2D antibody blockade to prevent ligand binding after IT Treg ablation, and observed significantly reduced control of MC38-*B2m^-/-^* tumors (Fig. 2J). This suggests that activating ligand binding by NKG2D (and likely other NK activating ligands) can support tumor control in this context. Next, we examined whether activating ligand binding could mediate tumor control even against MHC I+ tumors. We used MHC I+ RMA tumor lines that do not naturally express NKG2D ligands and then introduced the NKG2D ligand H60 to test the importance of NKG2D-mediated recognition for tumor control. Importantly, as NKG2D has been noted to impact CD8^+^ T cell killing and the H60a antigen is a foreign antigen in C57Bl6 mice, all mice in these experiments were depleted of CD8^+^ T cells to rule out any potential CD8^+^ T cell response to MHC I+ tumors in this setting^41,42^. MHC I+ RMA-WT tumors, which lack NKG2D ligands and are known to be resistant to NK cells^46^, were not controlled by IT Treg ablation (Fig. 2K). In contrast, when these cells were transduced to express the NKG2D ligand H60a (Ext. Data Fig. 2g), they were controlled by IT Treg ablation in an NK-dependent fashion (Fig. 2L), and NKG2D blockade significantly reduced tumor control (Fig. 2M). In a similar experiment using the MHC I+ B16F10 tumor model, which also does not express NKG2D ligands, we observed a similar tumor control that was NK cell- and NKG2D dependent when the tumor cells overexpressed a different NKG2D ligand, RAE-1b (B16F10-RAE-1b) (Ext. Data Fig. 2h). Therefore, IT Treg depletion can mobilize NK cell antitumor responses against both MHC I-high and MHC I-low tumor cells by engaging NKG2D ligands on tumor cells. Overall, we hypothesize that Treg ablation can induce NK cell control of MHC I-deficient tumors with little or no activating ligand expression, but can also elicit NK cell control of MHC I+ tumors if tumor cells express sufficient NK activating ligands to overcome inhibition via inhibitory receptors specific for MHC I. Thus, Treg ablation enhances NK cell control of tumors mediated by both missing-self recognition and "induced-self" recognition.

### IT Treg ablation elicits transcriptional changes in NK cells that depend on CD4^+^ Tconv cells

To investigate changes in NK cells using an unbiased approach, we performed bulk RNA sequencing analysis of NK cells (*DAPI^-^CD45^+^CD3^-^CD8a^-^CD19^-^F480^-^NK1.1^+^*) FACS sorted from MC38-*B2m^-/-^* tumors in *Foxp3^DTR-GFP^* mice six days after IT injections of PBS or DT, or DT with systemic depletion of all CD4^+^ T cells (Fig. 3A-B). PCA analysis showed a shift in the NK cell transcriptome associated with IT Treg depletion compared to the PBS control (Fig. 3C, Ext Data Fig. 3a), and volcano plot analysis showed increased expression of several genes associated with NK cell activation and cytotoxic status (Fig. 3D-E, Supplementary Table 1).

**Figure 3:**
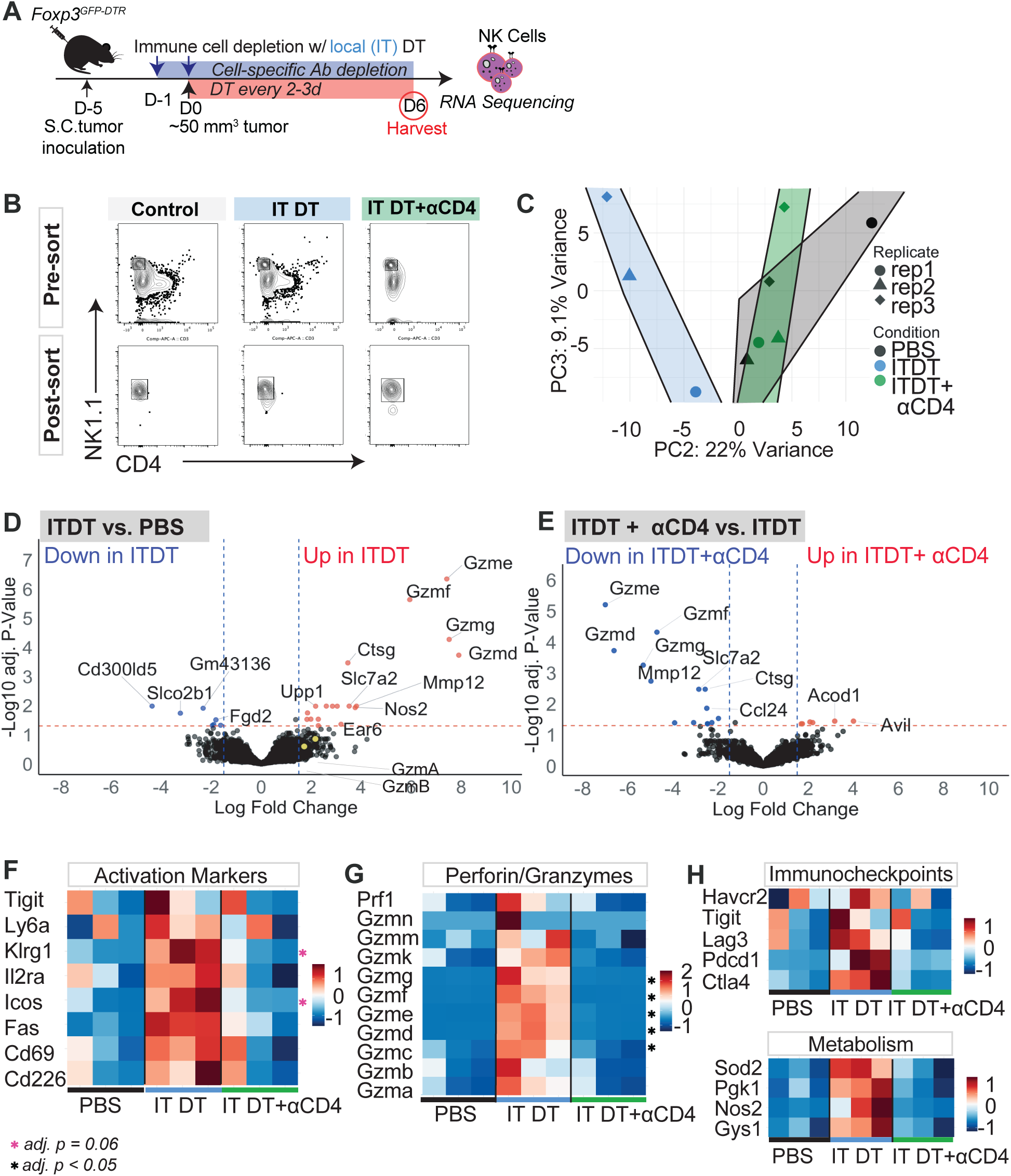
Global transcriptomic changes in NK cells following IT-Treg ablation or ablation of both Tregs and CD4 T cells. (A) Schematic of MC38-*B2m-/-* tumor inoculation and treatment regimen in *Foxp3^GFP-DTR^* mice depleted of Tregs intra-tumorally, before tumor resection 6 days later for downstream processing and sorting to obtain NK cells from three conditions: PBS, IT DT or IT DT+ anti-CD4. RNA sequencing was performed on three individual replicates, each comprised of sorted cells from a pool of 3-4 tumors. (B) Representative flow plots for pre- and post-sort NK cells from three conditions: PBS, IT DT or IT DT+ anti-CD4 (C) Principal component analysis (PCA) plot of PC2 vs PC3 showing clustering of NK cells with the various treatment conditions. In this analysis PC1 (not shown) mainly reflected differences in the replicates. (d-e) Differential expression analysis was conducted with limma (Methods). Volcano plots show upregulated and downregulated genes in NK cells in the three different conditions. Genes were selected as differentially upregulated (red) or downregulated (blue) and highlighted if adjusted p-value <0.05 and log 2 fold change >1.5 in absolute value. (f-h) Heatmaps of z-normalized expression for selected genes encoding activation markers, granzyme family members, immune checkpoint receptors (ICP), and metabolic markers. Genes marked with an asterisk (black) are significant and have adjusted p-values < 0.05, or marked with an asterisk (pink) if adjusted p-values = 0.06, based on differential analyses in (d-e).

We used the z-scaled rotation (=loadings) from the PCA and selected genes from different pathways to assess using a z-score transformation of transcripts per million (TPM). IT Treg ablation led to a prominent increase in the expression of genes encoding common NK cell activation markers, i.e. *Tigit, Ly6a* (SCA-1), *Klrg1, Icos, Fas, CD69* and *CD226* (DNAM-1), as well as numerous granzyme genes, and immune checkpoint (ICP) genes *Havcr2* (TIM3), *Tigit*, *Lag3*, *Pdcd1* and *Ctla4* (Figs. 3F-H). Surprisingly, when Tregs were ablated, orphan granzyme family members (*GzmC-N)* were more upregulated than conventional granzymes A and B (Fig 3D, G). Although poorly studied in both humans and mice, these other granzymes can play important roles in the immune system^43^. Key metabolic pathway genes were also upregulated in NK cells, including *Nos2* (nitric oxide synthesis pathway)^44^*, Gys1* (glycogen synthesis pathway)^45^*, Pgk1* (glucose metabolism and ATP production)^46^, and *Sod2* (mitochondrial reactive oxygen species metabolism)^47^, consistent with an oxidative and/or hypoxic stress response, metabolic reprogramming and a general inflammatory state (Fig. 3H).

Notably, the shifts in the NK cell transcriptome due to IT DT treatments were largely prevented by depleting all CD4^+^ T cells at the same time DT treatments were initiated (Fig. 3C-H, Ext. Data Fig. 3b^48^ Supplementary Table 2), suggesting that CD4^+^ Tconv cells were necessary for the altered transcriptome of NK cells upon IT Treg ablation. While CD4^+^ Tconv are recognized for their roles in priming, activating and providing key cytokines for CD8^+^ T effector cells^49–52^, less is documented concerning their roles in helping innate lymphocytes such as NK cells.

### IT Treg ablation increases NK cell activation, maturation, and cytotoxicity in a CD4 T cell-dependent manner

To correlate these transcriptomic findings to changes in protein levels and functional activities, we collected MC38-*B2m^-/-^* tumors at the same timepoint as the RNAseq data (day 6 after IT DT), as well as an earlier timepoint (day 4), and performed spectral flow cytometry analyses (Fig. 4A). At day 4, NK cells in the tumors from IT Treg- depleted mice were more proliferative, as demonstrated by *in vivo* labeling with BrdU (Fig. 4B), and expressed higher levels of the early activation marker CD69 (Fig. 4C). By day 6, additional activation markers for NK cells, SCA-1 and KLRG1, were also significantly upregulated, and NK cells exhibited a more mature phenotype (CD27^-^ CD11b^+^) (Fig. 4D-F). These phenotypic changes were abolished when CD4^+^ Tconv cells were co-depleted with IT Tregs, confirming that the activation and maturation of NK cells was dependent on CD4^+^ Tconv cells.

**Figure 4.**
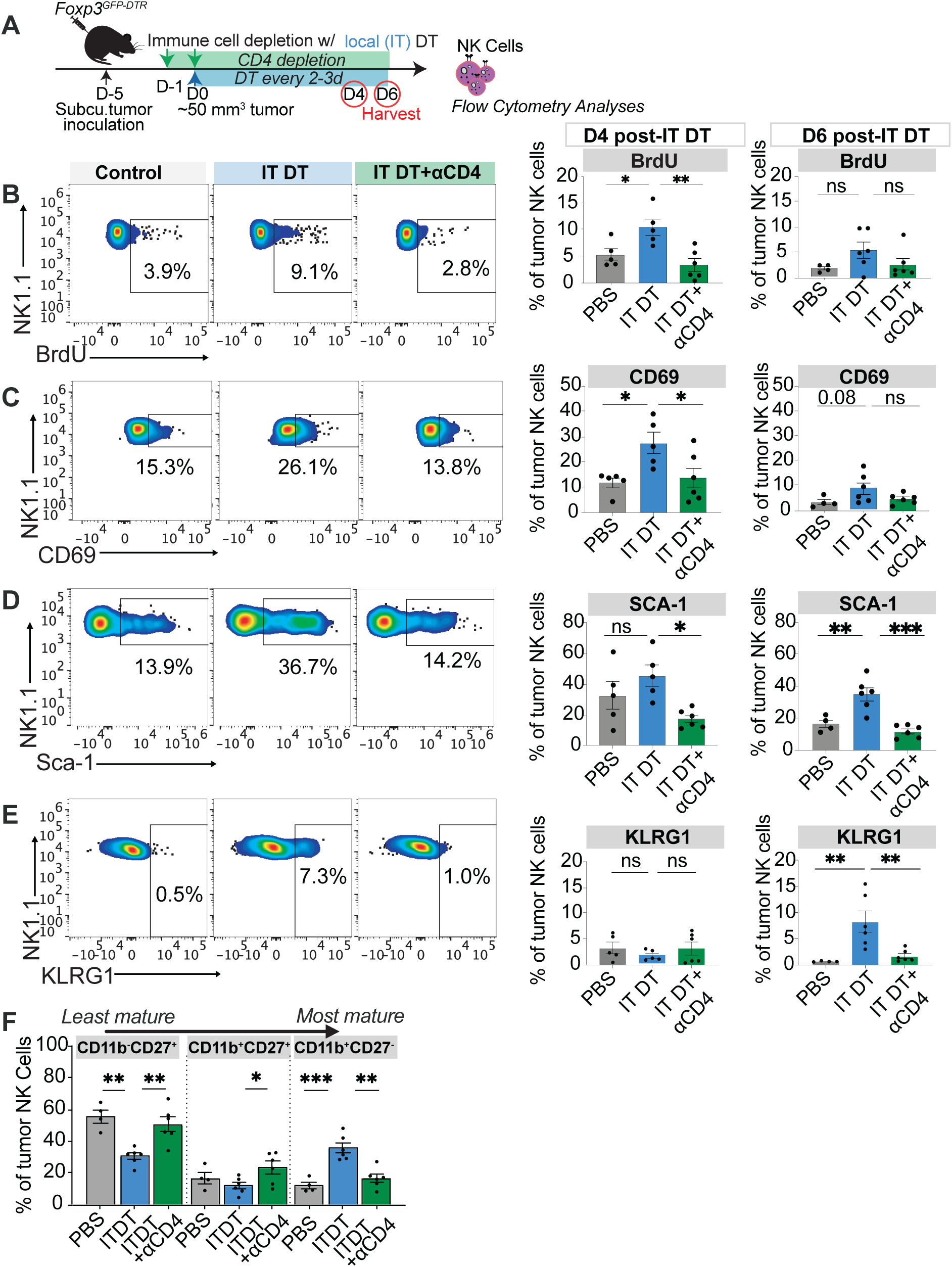
The NK cell activation phenotype following IT-Treg ablation is CD4 T cell-dependent. (A) Schematic of MC38-*B2m^-/-^* tumor inoculation in *Foxp3^DTR^* mice, IT depletion of Tregs, immune cell depletions, and tumor resection 4 or 6 days after treatment initiation, for phenotypic analysis of intratumoral NK cells. After processing, NK cells were sorted from tumors from three treatment conditions: PBS, IT DT and IT DT+ anti-CD4. (B-E) Representative flow plots (from day 4) and summary data (from days 4 and 6) of markers of proliferation (BrdU incorporation, B) and activation (CD69, C; Sca-1, D; and KLRG1, E) on gated NK cells in MC38-*B2m^-/-^* tumors from various treatment groups (PBS control in black, ITDT in blue, and ITDT + anti-CD4 in green). n=5-8 mice/group, representative of at least 3 independent experiments. (F) Percentages of NK cells with the indicated maturation phenotypes defined by CD11b and CD27 expression at D6 after starting IT treatments in MC38-*B2m^-/-^* tumors. The BrdU experiment was done twice and the other analyses were done 4 times. Data represent means ± SEM; *p < 0.05, **p < 0.01, ***p < 0.001, and ****p < 0.0001 from unpaired t-test comparisons. (n=5-8 mice/group). Ordinary one-way ANOVA with Dunnett’s multiple comparisons test.

Confirming the RNAseq data, IT Treg depletion also increased the intracellular levels of granzyme B and perforin (Fig. 5A-B). Importantly, *ex vivo* cytotoxicity of NK-sensitive MHC I-deficient tumor cells by sorted tumor NK cells was also elevated by Treg- depletion within tumors (Fig. 5C-D). Spleen cells from mice depleted of Tregs by systemic exposure to DT also showed elevated cytotoxicity *ex vivo*, which depended on the presence of CD4^+^ Tconv cells (Ext. Data Fig. 3c-e). The splenic effector cells were confirmed to be NK cells by both depleting and enriching them from these spleen cell populations *ex vivo* (Ext. Data Fig. 3d, e). Thus, Tregs prevented CD4 T cell-dependent induction of NK cell cytotoxicity in tumor-bearing mice.

**Figure 5.**
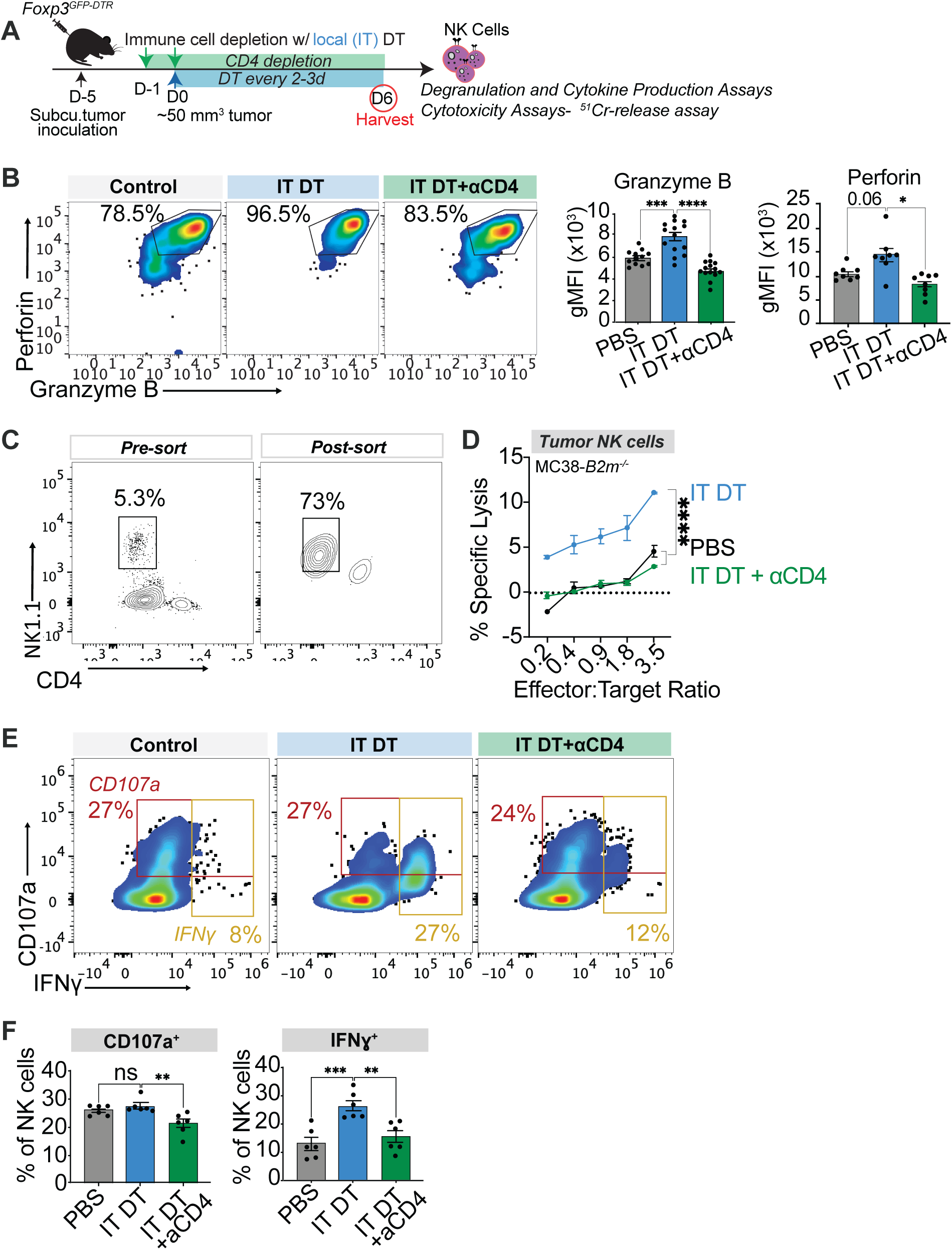
Induction of functional markers on NK cells following IT-Treg ablation is CD4 T cell-dependent. (A) Schematic of MC38-*B2m^-/-^* tumor inoculation and treatment regimen in *Foxp3^DTR^* mice to prepare NK cells from tumors for RNA and phenotypic analysis, from three treatment conditions: PBS, IT DT and IT DT+ anti-CD4. (B) Representative flow plots and quantification of MFI of intracellular granzyme B and perforin in NK cells in MC38-*B2m^-/-^* tumors of various treatment groups 6 days after initiation of DT treatments. Bar graphs are of data pooled from 2 independent experiments. (C, D) Cytotoxicity of MC38-*B2m^-/-^* target cells measured with 4-hour chromium release assays of sorted NK cells from MC38-*B2m^-/-^* tumors in mice treated with PBS, ITDT or ITDT+anti-CD4 (n=5-6 from each group). The target cell spontaneous release was 10.8% in this assay. (E, F) Representative flow plots and quantification of NK cell functional markers for degranulation (CD107a) and IFNg production across various treatment groups after stimulation with plate-bound NK1.1 antibodies ex vivo. Data represent means ± SEM; *p < 0.05, **p < 0.01, ***p < 0.001, and ****p < 0.0001 from Ordinary one-way ANOVA with Dunnett’s multiple comparisons tests. (n=5-8 mice/group, representative of at least 3 independent experiments).

Tumor NK cells from IT Treg-depleted mice also showed significantly increased IFN-γ production when stimulated *ex vivo* with plate-bound NK1.1 antibodies, which was again abolished when the mice were also depleted of CD4^+^ Tconv cells (Fig. 5E-F, Ext. Data Fig. 3f). In contrast, we observed no increase in *ex vivo* degranulation (CD107a) upon NK1.1 stimulation, despite the increased killing noted earlier. We propose that although the level of degranulation in NK cells was not noticeably increased, the increased granzymes and perforin in the granules of these NK cells after Treg depletion can account for the increased cytotoxicity they exhibit. Thus, NK cells in tumors that were Treg- depleted exhibited increased proliferation, maturation, and tumor cell cytotoxicity. In each case CD4^+^ Tconv cells were required to impart these changes in NK cell function.

### Conventional dendritic cells are required for NK cell-dependent tumor control

Tregs regulate the activation and priming of effector T cells directly and also indirectly by suppressing dendritic cells^24,53–55^. Specifically, conventional dendritic cells expand and mature after systemic depletion of Tregs, and cDC1s (CD8α^+^/CD103^+^) have been shown to be important for activating CD8^+^ T cell responses, while cDC2s (CD11b^+^/Sirpα^+^) play a dominant role in stimulating CD4^+^ T cell responses^56–61^. In addition, NK cells have been implicated in recruiting cDC1s to boost CD8^+^ T cell responses^62,63^, and the direct stimulation of NK cells by DCs has also been described (find citations). However, whether DC-NK cross-talk is altered by Treg ablation has not been explored^65–67^.

To probe the contribution of cDCs in the antitumor response after Treg depletion, we generated bone marrow chimeras (BMCs) in which donor bone marrow cells from *Foxp3^DTR-GFP^;Zbtb46^DTR^*mice (to co-deplete all cDCs and Tregs), or *Foxp3^DTR-GFP^;Batf3^KO^*mice (which lack cDC1s but retain cDC2s) or control *Foxp3^DTR-GFP^* mice were used to repopulate lethally irradiated B6 mice (Fig. 6A). BMCs were necessary for these studies because DT treatment of intact *Zbtb46^DTR^* mice is lethal^64^. After 6 weeks to allow immune cell reconstitution, tumors were established in the mice and once tumors were palpable, IT DT treatments were initiated to deplete Tregs alone or in combination with cDC depletions. In the *Foxp3^DTR-GFP^;Zbtb46^DTR^* BMCs, we confirmed that there were significant decreases in bulk CD11c^+^MHCII^+^ DCs, as well as specifically both cDC1s and cDC2s, following DT administration (Ext Data Fig. 4a-c), and this resulted in the failure to control tumors with IT Treg ablation (Fig. 6B-C). In contrast, *Foxp3^DTR-GFP^;Batf3^KO^* BMC controlled the tumors after Treg depletion nearly as well as *Foxp3^DTR-GFP^* mice. By deduction, these findings suggest that cDC2s and not cDC1s are required for tumor control following IT Treg ablation.

**Figure 6.**
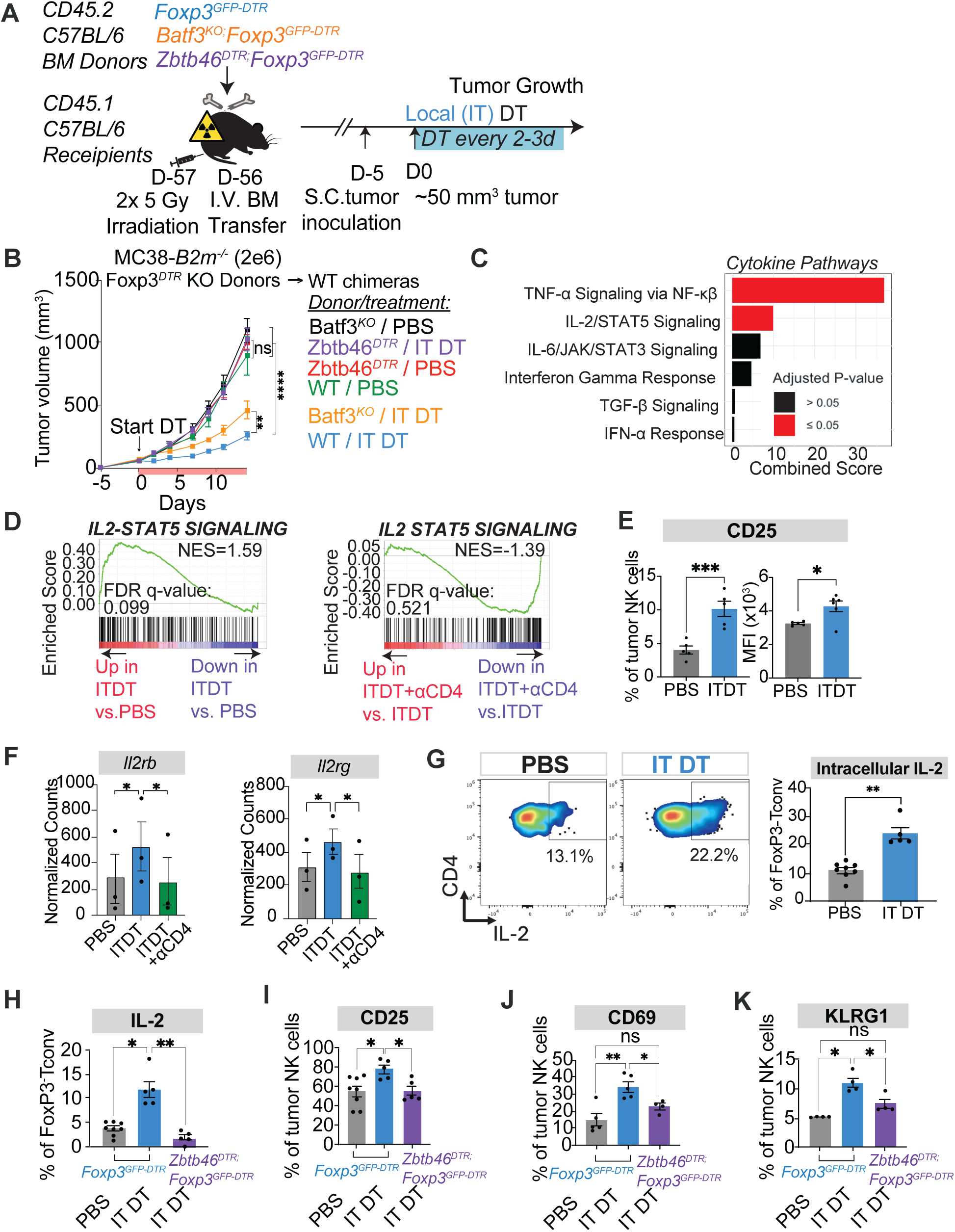
NK cell activation and anti-tumor activity is dependent on dendritic cells and IL-2 (A) Schematic of generation of bone marrow chimeras with *Foxp3^DTR^*, *Zbtb46^DTR^*;*Foxp3^DTR^* and *Batf3^-/-^*;*Foxp3^DTR^* hematopoietic cell compartments. After 8 weeks of reconstitution, MC38-*B2m^-/-^* tumor cells were inoculated and 5 days later the treatment regimen was initiated, to deplete Tregs or *Zbtb46* expressing cDC1/2s intra-tumorally. Tumors were resected at D6 after Tregdepletion for NK cell analysis. (B) Growth curves of MC38-*B2m^-/-^* tumors in mice that were IT depleted of Tregs (blue) or IT depleted of both Tregs and cDC1/2s (purple). Also tested were IT-Treg depleted mice lacking cDC1s (gold). Control groups in which Tregs were not depleted are shown for each mouse phenotype (green, black and red). The results shown are representative data from one experiment of three performed. (C) EnrichR highlighting cytokine signaling pathways for enrichment in genes enriched in ITDT versus PBS NK cells. (D) GSEA Enrichment plots for Hallmark IL-2/STAT5 signaling pathway using list of genes upregulated in ITDT versus PBS NK cells. FDR-q-value and normalized enrichment score (NES) are labeled in plot. (E) CD25 expression and MFI on tumor NK cells on day 6 following either PBS or IT- Treg ablation. (F) Transcripts per millions of *Il2rb* and *Il2rg* transcripts in NK cells from three conditions: PBS, IT DT or IT DT+ anti-CD4. Data are from the RNA sequencing dataset described in Fig 3. (G) Intracellular staining of IL-2 in tumoral CD4 Tconv cells (CD3+CD4+FOXP3-) 6 days after treating MC38-*B2m^-/-^* tumor-bearing mice with IT-DT (blue) or PBS (grey). Representative flow plots showing IL-2 staining are presented on the left with a summary on the right. Representative of 2 independent experiments. (H) Intracellular staining of IL-2 in CD4 Tconv cells (CD3+CD4+FOXP3-) on day 6 following either PBS, IT-Treg ablation, or IT-Treg and IT-cDC1/2 ablation in MC38- *B2m^-/-^* tumors. Representative flow plots are presented on the left, and summary data on the right. One experiment performed. (I) CD25 expression on tumor NK cells on day 6 following either PBS, IT-Treg ablation, or IT-Treg and IT-cDC1/2 ablation in MC38-*B2m^-/-^* tumors. One experiment performed. (J) CD69 expression on tumor NK cells on day 6 following either PBS, IT-Treg ablation, or IT-Treg and IT-cDC1/2 ablation in MC38-*B2m^-/-^* tumors. One experiment performed. (K) KLRG1 expression on tumor NK cells on day 6 following either PBS, IT-Treg ablation, or IT-Treg and IT-cDC1/2 ablation in MC38-*B2m^-/-^* tumors. One experiment performed. Data represent means ± SEM; *p < 0.05, **p < 0.01, ***p < 0.001, and ****p < 0.0001. B-C: RM two-way ANOVA followed by Geiser-Greenhouse correction. (n=5-8 mice/group, representative of at least 3 independent experiments). F, G, H, I, J: Ordinary one-way ANOVA with Dunnett’s multiple comparisons tests.

To interrogate how cDC2s and CD4^+^ T cells may contribute to NK cell activation, we determined whether Molecular Signatures Database (MSigDB) Hallmark datasets were enriched in NK cells following IT Treg depletion. Among the enriched pathways were multiple cytokine signaling pathways induced by cytokines that can be produced by T cells (Fig. 6D, Ext. Data Fig. 4d). Using Gene Set Enrichment Analyses (GSEA), we found that NK cells from Treg-depleted tumors were significantly enriched in genes associated with IL-2-STAT5 signaling and TNF-α signaling from the Gene Ontology (GO) Consortium Hallmark dataset (Fig. 6E, Ext. Data Fig. 4e, Supplementary Table 3). The enrichment in NK cells of genes associated with these datasets was largely dependent on the action of CD4^+^ Tconv cells (Fig. 6E, Ext. Data Fig. 4e). The interferon gamma response is commonly associated with effector functions and tumor immunity, but that pathway showed a low enrichment score that was not statistically significant (Fig. 6D, Ext. Data Fig. 4f).

Tregs suppress CD4^+^ T cell activation by suppressing their production of IL-2 and by acting as an IL-2 sink, among other mechanisms^65–69^. We found that cell surface IL- 2Rα (CD25) was increased on NK cells after IT Treg depletion (Fig. 6E), as were transcripts encoding the IL-2Rβ and IL-2Rγ subunits (Fig. 6F). To determine the source of IL-2 production following IT Treg ablation, we performed intracellular cytokine staining of immune cells from tumors by directly incubating the cells ex vivo in Golgi-blockers without *ex vivo* stimulation. This revealed that CD4^+^ Tconv cells were a prominent source of IL-2, and that IL-2 production increased in CD4^+^ Tconv cells with IT Treg depletion (Fig. 6G). Importantly, analysis of the aforementioned BMC mice lacking cDCs demonstrated that the increased IL-2 production from CD4^+^ Tconv cells after Treg depletion, as well as increased CD25, CD69 and KLRG1 expression on NK cells, were dependent on cDCs (Fig. 6H-K). Taken together, these data suggest that cDC2s are necessary to stimulate CD4^+^ Tconv cell production of IL-2 as well as for elevating IL-2R expression on NK cells.

### Anti-tumor NK cells require IL-2 from CD4^+^ Tconv cells

To test the importance of the cytokine signaling pathways enriched in NK cells after Treg depletion in promoting NK cell tumor control, we employed neutralizing antibodies against IFN-γ, TNF-α and IL-2 (Fig. 7A, Ext. Data Fig. 5a-d). In mice with either MC38-*B2m^-/-^* or B16-F10-*B2m^-/-^* tumors, IL-2 neutralization prevented tumor control following IT Treg ablation (Fig. 7B, Ext. Data Fig. 5a, d). In contrast, neutralization of IFN-γ or TNF-α did not prevent robust tumor control with IT Treg ablation (Ext. Data Fig. 5b-c). These data suggested that IL-2 signaling, but not TNF or IFN-γ signaling, was critical for tumor control in the context of IT Treg ablation. Furthermore, IL-2 neutralization prevented the induction of CD25, SCA-1, granzyme B, and acquisition of the mature CD27^-^CD11b^+^ phenotype by NK cells (Fig. 7C-F).

**Figure 7:**
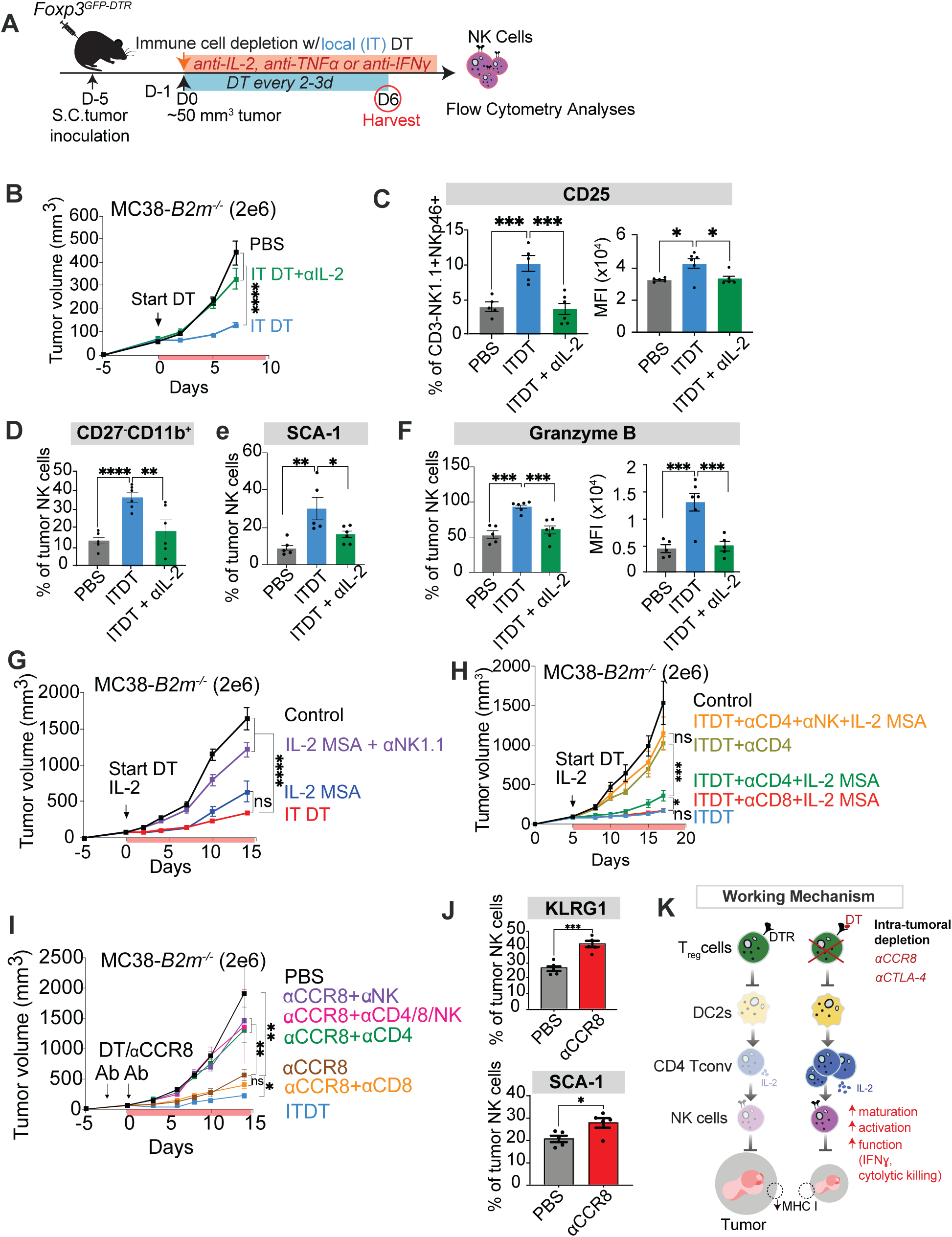
IL-2 signaling is essential for tumor control after IT-Treg ablation (A) Schematic of *MC38-B2m^-/-^* tumor inoculation and treatment regimen in Foxp3*^DTR^* mice. Tumors were resected at D6 after IT-DT for processing to obtain NK cells from various conditions for phenotypic analysis. (B) Growth curves of MC38-*B2m^-/-^* tumors in mice that were intra-tumorally depleted of Tregs (blue) in combination with anti-IL-2 (JES6-1) (green), or no Treg depletion (black). Representative of two independent experiments. B16F10-*B2m^-/-^* tumors show similar trend (not shown). (C) Percentages and MFI of CD25+ cells among tumoral NK cells in mice bearing MC38-*B2m^-/-^* tumors that were treated with DT IT 6 days previously, without (blue) or in combination with anti-IL-2 (JES6-1) treatments (green). Control (grey) mice were not Treg depleted. Representative of 3 independent experiments. (D-F) CD27-CD11b+ (mature phenotype), Sca-1 and Granzyme B staining of tumoral NK cells in the treatment groups described in (C). Percentages or mean fluorescence intensities are shown. Representative of four independent experiments. (G) Growth curves of MC38-*B2m^-/-^*tumors in mice with intact Tregs that were injected with IL-2-MSA or depleted of NK cells before treatment with IL-2-MSA, in comparison to no treatments (control). IT-Treg deleted mice are shown for comparison (red). Representative of 3 independent experiments. (H) Growth curves of MC38-*B2m^-/-^* tumors in IT-Treg depleted mice (blue), that were also depleted of CD4 cells (gold), or CD4-depleted and supplemented with IL-2-MSA (green), or the latter treatments combined with NK depletion (orange). Control mice in which Tregs were not depleted are shown for comparison (black). Representative of 3 independent experiments. (I) Comparing methods of Treg ablation with final tumor volumes at D15 post initiation of depletion in MC38-*B2m^-/-^* tumors. One experiment performed. (J) Growth curves of MC38-*B2m^-/-^* tumors in mice with treated with CCR8 antibody, combined with various cellular depletions. One representative experiment of three is shown. (K) Model of mechanism Data represent means ± SEM; *p < 0.05, **p < 0.01, ***p < 0.001, and ****p < 0.0001. B, G, H, J: RM two-way ANOVA followed by Geiser-Greenhouse correction. (n=5-8 mice/group). C-F: Ordinary one-way ANOVA with Dunnett’s multiple comparisons tests.

We hypothesized that IT Treg ablation resulted in the increased availability and production of IL-2 that activated the antitumor NK cell response. Therefore, we tested whether excess IL-2 was sufficient to drive tumor control even in the presence of Tregs. Administration of half-life extended IL-2, i.e. IL-2 fused to mouse serum albumin (IL-2- MSA)^34,36^, was sufficient to promote the control of MC38-*B2m^-/-^* tumors even when Tregs were present (Fig. 7G, Ext. Data Fig. 5e). NK cell depletion prevented tumor control elicited by IL-2-MSA administration, confirming that increased IL-2 mobilized NK cell-mediated antitumor activity (Fig. 7G, Ext. Data Fig. 5e).

To test whether IL-2 could replace the requirement for CD4^+^ Tconv cells in this setting, we supplemented IT-Treg depleted mice co-depleted of CD4^+^ T cells with IL-2- MSA. Indeed, exogenous supplementation of IL-2-MSA restored tumor control in a manner dependent on NK cells, but not CD8^+^ or CD4^+^ T cells (Fig. 7H, Ext. Data Fig. 5F). Together, these data, along with the evidence of an activated IL-2 signaling pathway in

NK cells, argue that after IT-Treg ablation, cDC2s stimulate CD4^+^ Tconv cells to produce IL- 2 that acts directly on NK cells to drive NK-mediated tumor control, highlighting an important axis of cell interactions unleashed by local IT Treg depletion.

### Clinically applicable intra-tumoral Treg depleting antibodies promote tumor control dependent on NK cells and CD4^+^ Tconv cells

Studies have demonstrated that certain systemic antibody treatments, including anti-CTLA4 and anti-CCR8, can selectively deplete IT Tregs without depleting Tregs systemically throughout the body^9,70,71^. To extend these findings to MHC I-deficient tumors, we assessed CCR8 expression on IT Tregs and confirmed earlier findings that it is highly elevated on Tregs infiltrating MHC I-deficient tumors compared to those in the spleen^72^ (Ext. Data Fig. 6a). Anti-CCR8 treatment significantly decreased IT Treg percentages without decreasing splenic Treg percentages (Ext. Data Fig. 6b). Most importantly, systemic administration of anti-CTLA4 or anti-CCR8 resulted in control of established MHC I-deficient tumors, comparable to the impact of IT DT treatments in *Foxp3^DTR-GFP^* mice (Fig. 7I, Ext. Data Fig. 6c-e). Similar to our findings with depletion of IT Tregs in *Foxp3^DTR-GFP^* mice, tumor control with anti-CCR8 antibodies depended on NK cells and CD4^+^ T cells, but not on CD8^+^ T cells (Fig. 7I). NK cells were also shown to increase expression of activation markers KLRG-1 and Sca-1 with anti-CCR8 (Fig. 7J). Therefore, clinically applicable strategies using systemically infused antibodies to selectively deplete IT Tregs can control tumors that lack MHC I by mobilizing CD4^+^ T cells and NK cells.

Based on our findings, it is clear that Tregs can regulate antitumor NK cell responses, especially against CD8-resistant, MHC I-deficient tumors. Our data revealed that intra-tumoral Treg depletion enables CD4^+^ Tconv cells to produce IL-2 in a manner that is dependent on cDC2s, which acts on NK cells to promote their cytotoxic function against tumor cells, thus highlighting an important NK-CD4-cDC2 axis of cellular interactions within the tumor microenvironment (Fig. 7K).

## DISCUSSION

In the present study, we demonstrated that depleting Tregs locally within tumors can control multiple tumor types even when deficient for MHC I. IT Treg ablation also elicited tumor control without any autoimmune pathologies that are associated with systemic Treg depletion. Control of MHC I-deficient tumors required NK cells and CD4^+^ Tconv cells. Importantly, local Treg depletion also sensitized MHC I+ tumors to NK cell killing if tumors expressed NKG2D activating ligands. *In vivo* cellular depletions combined with bulk RNA sequencing and phenotypic and functional analyses of tumor infiltrating NK cells *ex vivo* demonstrated that a major contribution of CD4^+^ Tconv cells to the NK cell response was their production of IL-2, which activated NK cell functions directly within tumors. The co-depletion of CD4^+^ Tconv cells with Tregs prevented NK cells from upregulating markers of activation and maturation, increasing expression of granzymes and perforin, and acquiring cytotoxic activity. Whereas conventional dendritic cells (cDCs) were essential for the NK cell-mediated antitumor response, cDC1s were dispensable, suggesting a crucial role for cDC2s. Thus, IT Treg depletion unleashed cDC2-mediated CD4^+^ Tconv cell activation and IL-2 production, which acts on NK cells to promote their maturation into cytotoxic killers of tumor cells.

Previously, the impact of regulatory T cell (Treg) depletion has been studied extensively in various contexts—in homeostasis^18,69^, autoimmune disease^73^, during viral infections^19^, and in the setting of cancer^16,29^. However, the focus has primarily been on Treg suppression of CD8^+^ T cell responses, or how suppression of CD4^+^ Tconv cells or APCs diminishes CD8^+^ T cell responses^74–76^. Limited studies have investigated the impact of Treg depletion on NK cell responses to tumors, and none have addressed the impact of local, IT Treg depletion on modulating the NK cell anti-tumor response. Because MHC I- deficient tumors that are resistant to CD8^+^ T cell recognition arise frequently, especially in the context of checkpoint immunotherapy of cancer, our findings reveal that Treg depletion can still have therapeutic importance by promoting NK cell responses that can kill MHC I-deficient cancer cells^56,57^.

Further, our approach of using local Treg depletion within tumors is promising as an immunotherapy given that it avoids lethal pathology and autoimmunity that have been shown to accompany systemic Treg ablation^18^. Systemically administered antibodies, such as anti-CTLA4 and anti-CCR8, can selectively deplete IT Tregs, and our results show that both treatments were effective against MHC I-deficient tumors, revealing a clinically translatable approach for effective Treg depletion therapies. While CTLA-4 blockade is complicated by its capacity to block inhibitory signaling in effector T cells, CCR8 antibodies appear to function more exclusively by depleting Tregs within tumors^77^. Importantly, local Treg ablation in tumors with IT DT or anti-CCR8 was not associated with detectable pathology. The efficacy of IT Treg depletion suggests that the response is elicited locally, but we cannot rule out the possibility that low-dose DT delivered IT may also reach adjacent sites, such as the draining lymph nodes, to exert its therapeutic effect.

Our data establish that a major function of conventional CD4^+^ T cells in the antitumor response to MHC I-deficient tumors is to provide IL-2 to NK cells. Indeed, we found that IL-2-MSA injections could replace the requirement of CD4^+^ Tconv cells for tumor control with IT Treg depletion. Thus, our data supports the hypothesis that Treg depletion activates NK cells due to increased IL-2 production from CD4^+^ Tconv cells and the removal of the Treg IL-2 sink^30^. The importance of IL-2 is in line with the well-known role that IL-2 plays in NK cell activation. Furthermore, we reported previously that an IL-2 superkine elicits NK-dependent therapeutic efficacy against MHC I-deficient tumor cells at a stage when NK cell desensitization was established in vivo^7,34^. Thus, the excess IL-2 associated with Treg depletion in our studies may promote tumor rejection in part by reversing or preventing NK cell desensitization. Although exogenous IL-2-MSA replaced CD4^+^ Tconv cells in our experiments, additional IL-2-independent mechanisms whereby CD4^+^ T cells promote NK cell responses may exist that are dispensable when excess IL-2 is available. Regardless, the results bolster the evidence that IL-2 or IL-2 superkines can exert potent antitumor effects alone, or in combination with other immunotherapies, against both MHC I-deficient and MHC I+ tumors^7,78^. Interestingly, we found that IL-2- MSA injections stimulated NK cell-dependent antitumor responses even in the presence of Tregs. While these data are consistent with the possibility that Tregs suppress NK cell responses solely by regulating IL-2 availability, we cannot rule out that Tregs utilize additional mechanisms to suppress NK cell responses. Thus, further investigation of whether Tregs directly suppress NK cell activation independently of CD4^+^ Tconv cells is warranted.

We observed an essential role of cDCs in the antitumor response following IT Treg ablation. The finding that cDC1s were not required suggested a requirement instead for cDC2s, which aligns with the established requirement for cDC2s in activating most CD4^+^ Tconv cells across various types of immune responses^61^. Specifically, it was previously shown that subsets of cDC2s can traffic from tumor to tdLN and present tumor-derived antigens to CD4^+^ Tconv, and that following Treg depletion, the frequencies of these cDC2s and their capacity to elicit strong CD4^+^ Tconv responses against tumors were increased^61,79^. What was not known was that cDC2s can also influence NK cell responses to tumors. Together, our results highlight a Treg-CD4^+^ Tconv-cDC2-NK cell axis in antitumor immunity (Fig. 7k), offering new perspectives on the complex interplay between different immune cell subsets in the tumor microenvironment beyond the well-described Treg-cDC1-CD8^+^ T cell axis.

In conclusion, our findings open new avenues for cancer immunotherapy by revealing the potential to mobilize NK cells and CD4^+^ T cells against CD8^+^ T cell-resistant cancers following localized, intra-tumoral depletion of Tregs. Because Treg depletion can also drive immune responses against MHC I+ tumors, this strategy may effectively complement current checkpoint blockade immunotherapies that drive tumor resistance to CD8^+^ T cell responses. The fact that IT Treg depletion was sufficient to drive tumor control without inciting systemic autoimmunity provides a strong foundation for developing more effective and broadly applicable immunotherapeutic strategies.

## MATERIALS AND METHODS

### *In vivo* animal studies

All mice used were between 8-12 weeks of age and were compared to littermates or age-matched control mice. *Foxp3^DTR-GFP^* mice, kindly provided by Dr A. Rudensky (Memorial Sloan Kettering Institute)^21^, express human diphtheria toxin receptor and EGFP genes at the *Foxp3* locus without disrupting endogenous expression of the *Foxp3* gene. NKDTA mice were generated by crossing *Ncr1^iCre^* mice to *Rosa26^LSL-DTA^* mice (Jackson Laboratories). In *Foxp3^DTR-GFP^*mice, Tregs were depleted intra-tumorally by injecting 250 ng DT i.t. in x 50 ul every other day, or depleted systemically by injecting 1 ug DT in x 200 ul every other day. Diphtheria toxin was prepared in sterile HBSS solution at 1000 ng/ml and aliquoted and stored in -80 deg until use.

### Cell lines

RMA-*B2m^-/-^*, B16F10-*B2m^-/-^* and MC38-*B2m^-/-^* tumor lines were previously described^35^. RMA is a lymphoma, MC38 is a colorectal cancer line and B16F10 is a melanoma cell line, all from B6 mice. RMA-*B2m^-/-^* cells were cultured in RPMI 1640 (ThermoFisher) medium, whereas B16-F10-*B2m-/-* and MC38-*B2m^-/-^* cells were cultured in DMEM (ThermoFisher), in both cases in 10% FBS (Omega Scientific), 0.2 mg/ml glutamine (Sigma-Aldrich), 100 U/ml penicillin (Thermo Fisher Scientific), 100 µg/ml streptomycin (Thermo Fisher Scientific), 10 µg/ml gentamycin sulfate (Lonza), 50 µM β-mercaptoethanol (EMD Biosciences), and 20 mM HEPES (Thermo Fisher Scientific, in 5% CO2. All cell lines tested negative for mycoplasma contamination.

### *In vivo* tumor transplant experiments

Syngeneic C57BL/6J mice were inoculated with 2x10^6^ MC38-*B2m^-/-^* cells, 1x10^6^ B16-F10- *B2m^-/-^* cells or 1x10^6^ RMA-*B2m^-/-^* cells, in 100 uL of RPMI medium without serum. Tumor measurements were performed blindly across the entire experiment by a single operator with electronic calipers three times a week. Tumor volume was estimated using the ellipsoid formula: V = (4/3)πabc where a, b and c correspond to height, width and length of the tumors. Five days after tumor cell inoculation, when tumors reached volumes of 50 mm^3^, they were injected intra-tumorally (IT) with PBS or DT. Tumor experiments were terminated once the average of the length and width of tumors exceeded 20 mm. All experiments were conducted according to the Institutional Animal Care and Use Committee guidelines of the University of California, Berkeley.

### *In vivo* cellular depletions and cytokine blockade

To deplete NK cells, CD8 cells and/or CD4 cells, mice were injected IP with 200 µg of depleting antibodies specific for NK1.1 (clone PK136, Leinco), CD8β.2 (clone 53-5.8, Leinco) or CD4 (clone GK1.5, Leinco), respectively, 1 day before and on the day of initiating Treg depletions, and again every six days thereafter until mice were euthanized or no palpable tumors were detected for one week. Anti-CTLA4 (clone 9D9) and anti- CCR8 (GS-1811, Gilead Sciences) were dosed at 200 µg IP every 3 days. Cellular depletions were confirmed by flow cytometry. To neutralize cytokines, mice were injected IP with 200 ug doses of neutralizing antibodies specific for IL-2 (clone JES6-1) TNF-α (clone TN3-19.12 ) or IFN-γ (cloneXMG1.2), or control whole rat IgG (Jackson ImmunoResearch) starting on the day of Treg depletion and repeating every 3 days.

### Generation of bone marrow chimeras

Host CD45.1 C57/BL6 mice were irradiated using an Xrad320 (Precision X-Ray Irradiation) with two doses of 5 Gy, 16h apart. Four hours after the second dose of irradiation, mice received 8-10 million bone marrow cells via tail vein injection. Bone marrow cells were prepared by surgically resecting femurs and tibias from euthanized mice, which were then cleaned to remove excess muscle tissue, and bone marrows were flushed into a 15 mL conical tube using 27-gauge syringes containing ice-cold complete RPMI. Red blood cells were removed using ACK lysis buffer and the white cells were washed three times with PBS before being resuspended in serum-free RPMI and injected into mice via the tail vein. Following the bone marrow injections, mice received water supplemented with sulfamethoxazole-trimethoprim oral suspension (Ani Pharmaceuticals) for 30 days with daily monitoring.

### Flow cytometry

Tumors were resected from animals and diced into small fragments with a razor blade, followed by enzymatic digestion in RPMI medium containing 125 U/mL Collagenase D (Roche Diagnostics) and 20 mg/mL DNase I (ThermoFisher Scientific) using an orbital shaker for 45 min at 37C. Cells were stained directly for determining NK cell and T cell numbers and phenotypes, or incubated for 4 hours in medium containing Brefeldin A (Biolegend) and Monensin (Biolegend) for intracellular staining of granzyme B and perforin. LIVE/DEAD Fixablue blue or aqua stain (Molecular Probes) was used to exclude dead cells. Before staining with antibodies, FcγRII/III receptors were blocked by resuspending the cells in 50 µl undiluted culture supernatant of the 2.4G2 hybridoma cell line (prepared in the lab) and incubating for 10 minutes at 4C. Cells were washed in PBS containing 2.5% FCS and stained with antibodies directly conjugated to fluorochromes for 1h at 4C in the same buffer. For intracellular staining of granzyme B and perforin, cells were fixed and permeabilized using the FoxP3/ Transcription Factor Staining Buffer Kit (Tonbo) and stained with antibodies directly conjugated to fluorochromes for 1hr at RT in 1X Perm/Wash buffer (BD Biosciences). For tetramer staining, cells were incubated with the relevant PE-conjugated CD1d tetramers loaded with α-galactosylceramide or unloaded controls (NIH Tetramer Core Facility contract number 75N93020D00005) for 30 min at 4C, washed and resuspended before subsequent costaining. NK cells were gated as viable, CD45+, CD3-, CD19-, F4/80-, Ter119-, NK1.1+, NKp46+ cells. Flow cytometry was performed using an LSRFortessa, LSRFortessa X-20 (BD Biosciences) or Cytek Aurora Spectral Cytometer. Data were analyzed using FlowJo software (Tree Star).

### RNA Sequencing

Tumors from mice treated with PBS, IT-DT or IT-DT + anti-CD4 were resected, and single cell suspensions were prepared as described above. Tumor immune cells were enriched by layering on a lympholyte-M gradient (Cedarlane Laboratories Ltd., Canada) then stained with DAPI, CD19-PECy5, F4/80-PECy5, Ter119-PECy5, CD8a-PECy5, CD3e-APC, CD45-FITC, CD4-BV605, and NK1.1-APC-Cy7 for 30 min on ice. NK cells and CD4 T cells were sorted from tumors pooled from 2-4 mice per replicate, with a yield ranging from 20,000-200,000 cells, with three replicates total. To prepare RNA, cells were lysed in RLT buffer (Qiagen) containing 1% 2-mercaptoethanol (Thermo Fisher Scientific) before being purified with RNeasy Miniprep kits (Qiagen) and validated with Qubit, bioanalyzer and fragment analyzer (QB3 Genomics, UC Berkeley, Berkeley, CA, RRID:SCR_022170). Sequencing libraries were prepared using KAPA Hyper Prep Kit (KAPA Biosystems) and sent for bulk RNA sequencing with NovaSeq 6000 150PE (QB3 Genomics, UC Berkeley, Berkeley, CA, RRID:SCR_022170). Samples were processed and normalized to account for sequencing depth differences using Kallisto.

### Sequencing Analyses

Heatmaps: Z-scores were computed for each gene using the formula Z=(X−µ)/σ, where X is the gene’s expression count, µ is the mean, and σ is the standard deviation across samples. This standardization was performed using the scale function in R, with any missing values handled by imputation. The count matrix was transformed from wide to long format using pivot_longer, which allowed for easier plotting of gene expression data. The Z-scores were visualized using the ggplot2 package, with a color gradient where blue indicates low expression, white for average, and red for high expression. The assumptions of normal distribution inherent in Z-score calculations were acknowledged, along with the limitations regarding the sensitivity to outliers.

Differential Gene Expression: Gene expression was modeled using the formula *Y*_*i*_ ∼ 0 + *T*, where *Y*_*i*_ is the log-CPM value for gene *i*, *T* is the treatment (i.e. PBS, IT-DT, or IT-DT+aCD4). We then performed three pairwise comparisons (PBS vs IT-DT, PBS vs IT- DT+aCD4 and IT-DT vs IT-DT+aCD4) between the three treatment groups to identify genes with significant log2 foldchange >1 and adjusted p-value <0.05). This superset of significantly changing genes was used in downstream analyses. We used enrichR and GSEA for pathway enrichment analysis of the significant genes. For plotting gene quantifications, we transformed the count data using rlog (regularized logarithm) function ><0.05. Plots were generated using ggplot2 package.

### *In vivo* BrdU Labeling

To labeling proliferating cells, mice were injected IP with 1 mg of BrdU in 100 µL sterile 1X DPBS and sacrificed 18-24h later. Cells were stained with surface markers as described earlier. Cells were then fixed and permeabilized using the BD Cytofix/Cytoperm kit according to the manufacturer’s instructions. Intracellular BrdU staining was performed using anti-BrdU antibody (clone Bu20a, BioLegend) for 30 minutes at room temperature.

### *Ex vivo* cytotoxicity assay

In all cases cytotoxicity was assessed with a standard 4-hour ^51^Cr-release assay as described^35^, using NK sensitive MC38-*B2m^-/-^* tumor cells as target cells^35^. NK cells from tumors were sorted by flow cytometry from single cell suspensions of tumor cells as *DAPI^-^CD45^+^CD3^-^CD8a^-^CD19^-^F480^-^NK1.1^+^* cells. For testing cytotoxicity by splenocytes, NK cells were in some cases enriched from splenocytes using MojoSort^TM^ Mouse NK cell isolation kit, or depleted from splenocytes with biotin-anti-NK1.1 antibodies using the EasySep™ Mouse Strepavidin RapidSpheres™ Isolation Kit.

### *Ex vivo* stimulation assay

For assessing NK degranulation and cytokine production, single cell suspensions of splenocytes were incubated for 4 hours in high-binding 96-well plates pre-coated overnight at 4 deg with antibodies for the NK1.1 activating receptor (5 ug/ml), or isotype control antibodies (5 ug/ml), in DMEM medium containing Alexa Fluor-conjugated CD107a antibodies, 2.4G2 hybridoma supernatant to block FcγRII/III receptors, brefeldin A (Biolegend), and monensin (Biolegend) before performing surface and intracellular staining followed by flow cytometric analysis, gating on CD3-NK1.1+ cells.

| REAGENT or RESOURCE | SOURCE | IDENTIFIER |
| --- | --- | --- |
| **Antibodies** |  |  |
| Anti-CD45 BUV-395, clone 30-F11 | BD | Cat#: 564279; RRID:AB_2651134 |
| Anti-CD4 BUV737, clone RMA4-5 | BD | Cat#: 612843; RRID:AB_2870165 |
| Anti-CD4 APC, clone RMA4-5 | Biolegend | Cat#: 100516; RRID:AB_312719 |
| Anti-CD8a BV650, clone 53-6.7 | Biolegend | Cat#: 100742 |
| Anti-CD25 BV785, clone PC61 | Biolegend | Cat#: 102051; RRID:AB_2564131 |
| Anti-CD25 PerCPCy5.5, clone PC61 | Biolegend | Cat#: 102030; RRID:AB_893288 |
| Anti-CD25 BV711, clone PC61 | Biolegend | Cat#: 102049; RRID:AB_2564130 |
| Anti-CD44 APCe780, clone IM7 | eBioscience/Thermo Fisher | Cat#: 47-0441-82; RRID:AB_1272244 |
| Anti-PD1 BV711, clone 29F.1A12 | Biolegend | Cat#: 135231; RRID:AB_2566158 |
| Anti-CD103 BV785, clone 2E7 | Biolegend | Cat#: 121439 |
| Anti-CD69 PECy7, clone H1.2F3 | BioLegend | Cat#: 104512; RID:AB_493564 |
| Anti-CD62L Pacific Blue, clone MEL1-14 | Biolegend | Cat#: 104424; RRID:AB_493380 |
| Anti-IL10 APC, clone JES5-16E3 | Biolegend | Cat#: 505010; RRID:AB_315364 |
| Anti-TNFα BV510, clone MP6-XT22 | Biolegend | Cat#: 506339; RRID:AB_2563127 |
| Anti-IFNγ Pacific Blue, clone XMG1.2 | Biolegend | Cat#: 505818; RRID:AB_893526 |
| Anti-Foxp3 FITC, clone FJK-16S | Thermo Scientific | Cat#: 11-5773-82; RRID:AB_465243 |
| TruStain FcX™ PLUS (anti-mouse CD16/32) | Biolegend | Cat#: 156604; RRID:AB_2783138 |
| Anti-CD3e APC/fire 750, clone 17A2 | Biolegend | Cat #100248 |
| Anti-CD3e BV650, clone 17A2 | Biolegend | Cat #100229 |
| Anti-CD11b BV570, clone M1/70 | Biolegend | Cat # 101233 |
| Anti-CD27 BV650, clone LG.3A10 | Biolegend | Cat # 124233 |
| Anti-NK1.1 BV711, clone PK136 | Biolegend | Cat #108745 |
| Anti-NKp46 PE-Cy7, clone 29A1.4 | Biolegend | Cat #137618 |
| Anti-NKp46 PE, clone 29A1.4 | Biolegend | Cat #137604 |
| Anti-Ly6A/E (Sca-1) BV421, clone D7 | Biolegend | Cat # 108127 |
| Anti-Ly6A/E (Sca-1) BV785, clone D7 | Biolegend | Cat # 108139 |
| Anti-CD279 (PD-1) BUV737 | Biolegend | Cat #568363 |
| Anti-KLRG1 BV510, Clone 2F1 | Biolegend | Cat # 138421 |
| Anti-Granzyme A PerCP-eFluor 710, clone GzA-3G8.5) | Thermo Scientific | Cat #46-5831-82 |
| Anti-Granzyme B PE-Dazzle594, clone QA16A02 | Biolegend | Cat# 372216 |
| Anti-Perforin APC, clone S16009A | Biolegend | Cat# 154304 |
| Anti-EOMES PerCP-eFluor 710, clone Dan11mag | Thermo Scientific | Cat #46-4875-82 |
| Anti-CD19 PE-Cy5, clone 6D5 | Biolegend | Cat# 115510 |
| Anti-Ter119 PE-Cy5, clone TER119 | Biolegend | Cat# 116210 |
| Anti-F4/80 PE-Cy5, clone (BM8) | Biolegend | Cat# 123112 |
| Anti-T-Bet BV711 | Biolegend | Cat # 644820 |
| Anti-XCR1 AF647 | Biolegend | Cat #148214 |
| Anti-33D1 PE | Biolegend | Cat #124905 |
| Anti-CD11c BV785, clone N418 | Biolegend | Cat #117336 |
| Anti-Ly6C BV510 | Biolegend | Cat #128033 |
| Anti-H2K<sup>b</sup>/D<sup>b</sup> (MHC I) | Biolegend | Cat #114614 |
| Anti-IA/IE (MHC II) PE-Cy7 | Biolegend | Cat #107630 |
| Anti-CD107a (LAMP-1) AF647 | Biolegend | Cat #121610 |
| ***In vivo antibodies*** |  |  |
| Anti-NK1.1(PK136) | Leinco Technologies, Inc. | Lot # 1124L455 |
| Anti-CD4 (GK.15) | Leinco Technologies, Inc. | Lot # 0522L215 |
| Anti-CD8b.2 (53-5.8) | Leinco Technologies, Inc. | Lot # 1024L530 |
| Anti-IFNγ (XMG1.2) | Leinco Technologies, Inc. | Lot # 0818L380 |
| Anti-TNFα (TN3-19.12) | Leinco Technologies, Inc. | Lot # 0421L405 |
| Anti-IL-2 (JES6-1A12) | BioXCell | Cat # BE0043 |
| Anti-CTLA-4 (9D9) | BioXCell | Cat # BE0164 |
| Anti-NKG2D (MI-6) | eBioscience | Lot # E06727-1633 |
| Anti-CCR8 (GS-1811) | Gilead Sciences |  |
| **Chemicals, peptides, and recombinant proteins** |  |  |
| DNase I | Roche | Cat#: 10104159001 |
| Collagenase IV | Roche | Cat#: 11088882001 |
| Tonbo Foxp3/Transcription Factor Staining Buffer Kit | Tonbo Bioscience | Cat#: TNB-0607-KIT |
| Paraformaldehyde | Electron Microscopy Sciences | Cat#: 15713S |
| BrdU | BD Pharmingen™ | Cat # 51-2420KC |
| Triton X-100 | Fisher Scientific | Cat#: BP151-100 |
| Recombinant human IL-2 Teceleukin (Tecin™) | Hoffmann-LAROCHE | NCI repository (Frederick National Laboratory for Cancer Research) |
| IL-2-MSA | Produced in the laboratory of C. K. Garcia |  |
| BD GolgiPlug™ Protein Transport Inhibitor (Containing Brefeldin A) | BD Biosciences | Cat#: 555029 |
| eBioscience™ Fixable Viability Dye eFluor™ 780 | eBioscience | Cat#: 65-0865-14 |
| LIVE/DEAD™ Fixable Blue Dead Cell Stain Kit | Molecular Probes | Cat#: L34962 |
| LIVE/DEAD™ Fixable Aqua Dead Cell Stain Kit | Molecular Probes | Cat# L34965 |
| Percoll® Density Gradient Media | GE Healthcare | Cat#: 89428-522 |
| Diphtheria Toxin | Sigma-Aldrich | Cat # 322326 |
| CD1d OCH PE-Labeled Tetramer | NIH Tetramer Core Facility |  |
| CD1d Unloaded PE-Labeled Tetramer (control) | NIH Tetramer Core Facility |  |
| **Critical commercial assays** |  |  |
| EasySep™ Mouse CD4+ T cell Isolation Kit | STEMCELL Technologies | Cat#: 19852A |
| Dynabeads™ Mouse T-Activator CD3/CD28 for T cell Expansion and Activation | Gibco | Cat#: 11456D |
| NEBNext® Ultra II DNA library prep kit for Illumina® | New England Biolabs | Cat#: E7645S |
| NEBNext® Multiplex Oligos for Illumina® (Index Primer set 1–3) | New England Biolabs | Cat#: E7710S, E7335S, E7335S |
| RNAeasy Micro kit | Qiagen | Cat#: 74004 |
| KAPA mRNA HyperPrep Kit | Roche | Cat#: KK8580 |
| MojoSort™ Mouse NK cell Isolation Kit | BioLegend | Cat#: 480050 |
| EasySep™ Mouse Streptavidin RapidSpheres™ Isolation Kit | STEMCELL Technologies | Cat#: 19860A |
| APC BrdU Flow Kit | BD Pharmingen™ | Cat #: 51-9000019AK |

## Supplementary Materials

Supplementary Table 1: Upregulated and downregulated genes in tumor NK cells following IT Treg ablation (accompanying volcano plot).

Supplementary Table 2: CD4 T cell-dependence in NK cell genes regulated by IT Treg ablation.

Supplementary Table 3: Enriched pathways analysis on tumor NK cells.

## Acknowledgments

We thank L. Zhang and E. Seidel for laboratory support, H. Sudholz, A. Karampatzakis for providing comments on the manuscript, C. Newsom-Stewart for comments on graphic design, and the members of the Raulet and DuPage laboratories for helpful discussions. We also thank Kartoosh Heydari, Melaine Delcroix, and Harman Dhaliwal of the UC Berkeley Cancer Research Laboratory Flow Cytometry. We thank the NIH Tetramer Core Facility (contract number 75N93020D00005) for providing CD1d OCH PE-Labeled tetramers. We also thank Gilead Science. Inc. for providing both funding and anti-CCR8 antibodies for the study.

## Funding

NIH R01-CA-270790 (D.H.R.) 1DP2CA247830-01 (M.D.)

Pew-Stewart Scholar and St. Baldrick’s Scholar with generous support from Hope with Hazel (M.D.)

HHMI (K.C.G)

NIH-RO1-AI51321 (K.C.G)

Predoctoral fellowships from the National Science Foundation (DGE 1752814) (CZ) University of California Cancer Research Coordinating Committee (CRCC) (CZ) UC, Berkeley Center for Research and Education on Aging (CREA) (K.S.) Postdoctoral CIRM Training Program (EDUC4-12790) (Y.J.)

Cancer Research Institute / Amgen Irvington Postdoctoral Fellowship (CRI3984) (Y.J.)

## Author contributions

Conceptualization and methodology: D.H.R, M.D., A.W. and C.Z.

Investigation and experiments: C.Z., C.C., A.B., E.J, K.S., Y.J, S.L., S.S., E.A., A.M., L.Z. Writing – original draft: C.Z., M.D. and D.H.R

Writing – review & editing: C.Z., M.D. and D.H.R Funding acquisition, D.H.R. and M.D.

Provided critical reagent: C.G.

## Competing interests

D.H.R. is a cofounder of Dragonfly Therapeutics and serves on their scientific advisory board. He also serves on the SABs of Vivere Oncotherapies and Lightcast Discovery. He served in the past on the scientific advisory boards of Aduro Biotech, Innate Pharma, and Ignite Immunotherapy. Gilead Science Inc provided CCR8 antibody and funding to M.D. for part of this work.

## Data and Materials Availability

### Lead Contact

Further information and requests for resources and reagents should be directed to and will be fulfilled by the lead contacts, David Raulet and Michel DuPage.

### Materials Availability

Mouse lines and tumor cell lines generated from this study are available from UC Berkeley.

### Data and Code Availability

The RNA-seq data generated in the study are available on the GEO database (GSE287581).

To review GEO accession GSE287581:

Go to https://www.ncbi.nlm.nih.gov/geo/query/acc.cgi?acc=GSE287581 Enter token <u>uvgfomwqrjktnsv</u> into the box.

R code will be made available by lead contact.

---

**Ext. Data Figure 1.**
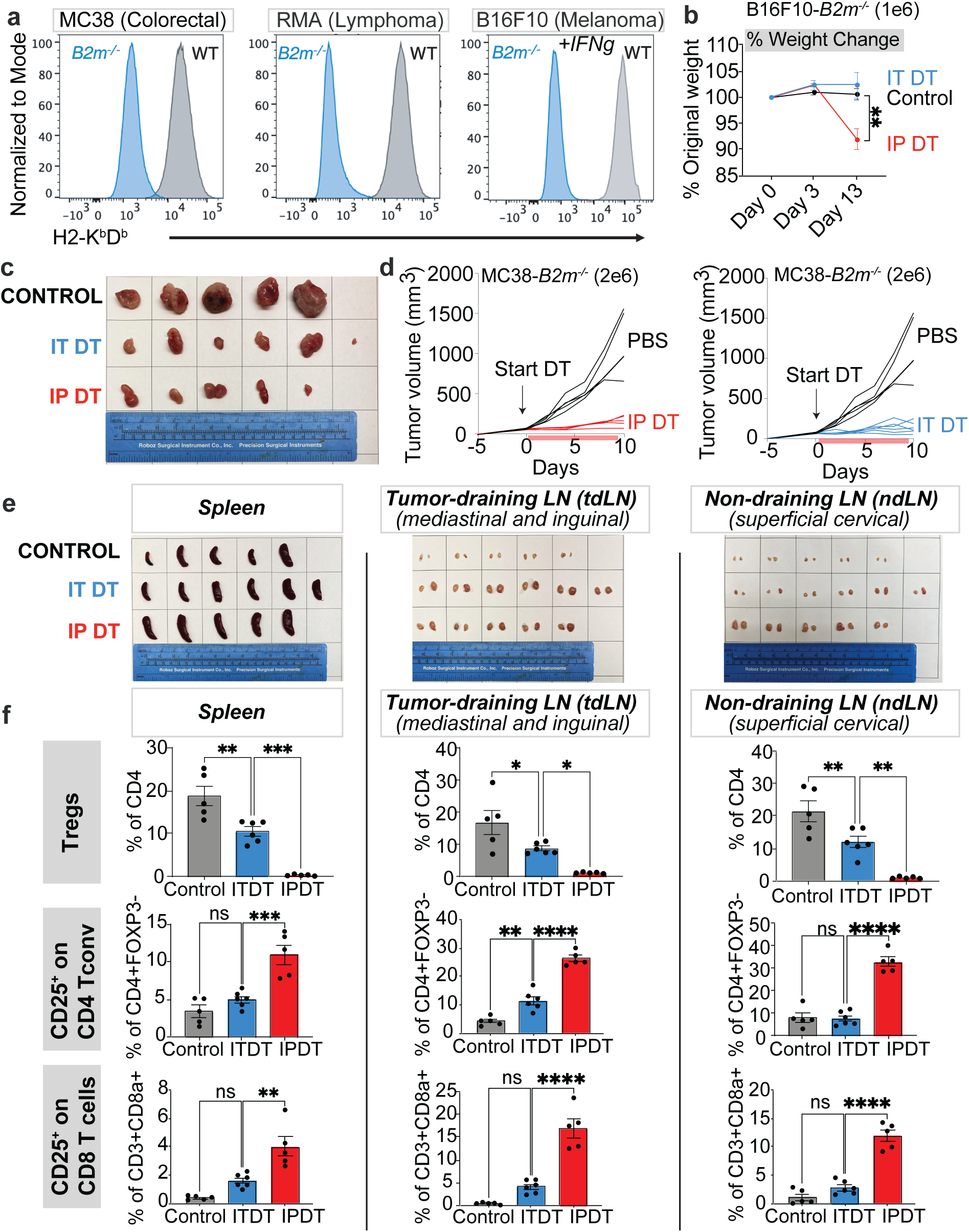
MHC I-deficient tumor lines and comparison of systemic vs. intra-tumoral Treg depletion models (a) Histograms showing staining of H2-K^b^/D^b^ antigens on WT (MHC I-expressing, grey) vs *B2m-/-* cell lines (blue). The B16F10 cells were stimulated overnight in vitro with 20 ng/mL IFNg to induce MHC I expression. (b) Body weight changes in *Foxp3^DTR-GFP^* mice with B16F10-*B2m^-/-^* tumors that were depleted of Tregs with IT vs IP DT injections. (c) Representative experiment showing relative MC38-*B2m^-/-^* tumor sizes at 12 days post Treg ablation (D17 post-tumor inoculation). (d) Spider plots for *Foxp3^DTR-GFP^* mice with MC38-*B2m^-/-^* tumors that were depleted of Tregs with IT vs IP DT injections. (e) Images of spleens, tdLN and ndLN from the experiment in panel (C). (f) Percentages of Tregs and CD25+ cells among CD4 Tconv cells (CD4+CD3+FOXP3) and CD8+CD3+ T cells from the spleens, tumor draining LN (tdLN), and nondraining LN (ndLN) 12 days after DT initiation. Data represent means ± SEMs; *p < 0.05, **p < 0.01, ***p < 0.001, and ****p < 0.0001 from Ordinary one-way ANOVA with Tukey’s multiple comparisons test, representative of at least 2 independent experiments.

**Ext. Data Figure 2.**
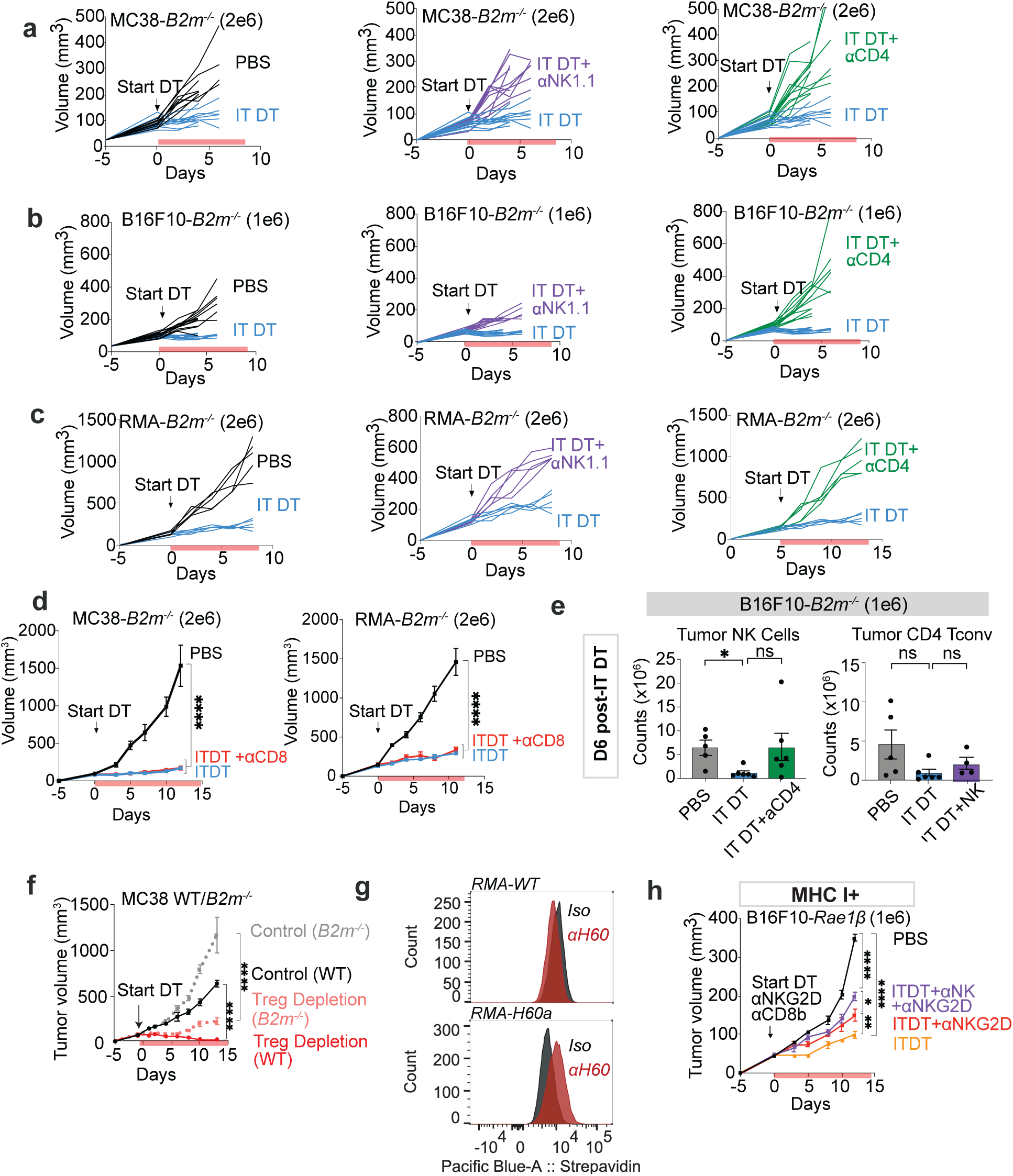
(associated with Fig 2). Cellular requirements for tumor rejection (a, b, c) Spider plots from Figure 2E, 2F and 2G (d) Growth curves of MC38-*B2m^-/-^* and RMA-*B2m^-/-^* in mice that were either treated with IT PBS, IT DT or depleted of CD8b+ T cells with IT DT. (e) Counts of NK cells and CD4 Tconv cells per tumor B16F10-*B2m^-/-^* tumors with the indicated treatments. (f) Growth curves of MC38-*B2m^-/-^* and MC38-*WT* tumors in mice that were intra- tumorally depleted of Tregs. Representative data from one of two experiments. (g) Staining validation for ectopic expression of NKG2D ligand H60a on RMA lymphoma tumor cell lines (RMA-WT vs RMA-H60a) using biotinylated polyclonal H60 antibodies. (h) Growth curves of B16-RAE-1b-transduced B16F10 tumors in mice that were depleted of CD8b+ T cells before starting IT DT and anti-NKG2D (clone MI-6) treatments. One experiment was performed. Data represent means ± SEM; *p < 0.05, **p < 0.01, ***p < 0.001, and ****p < 0.0001. d, f: Ordinary one-way ANOVA with Dunnett’s multiple comparisons test, representative of at least 1independent experiment per tumor cell line). h: RM two-way ANOVA followed by Geiser-Greenhouse correction.

**Ext. Data Figure 2.**
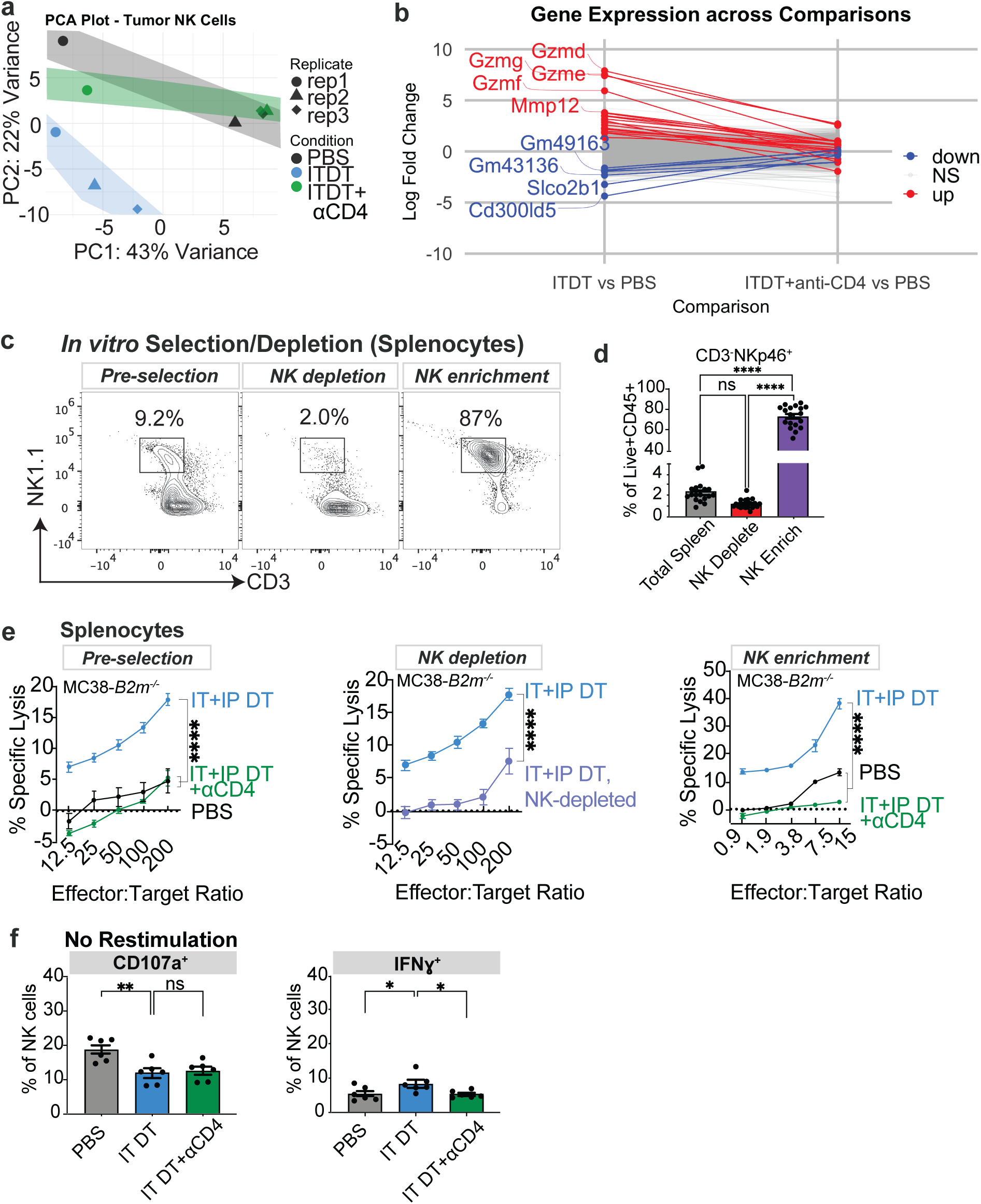
(associated with Fig 4, 5). Analyses of NK cells, ILC1 cells and invariant NK1.1+ T in tumors and spleens of treated mice. (a) Principal component analysis (PCA) plot of PC1 vs PC2 showing clustering of NK cells with the various treatment conditions (b) Volcano plots with dot and line overlaps^49^ of logFCs of ITDT_aCD4vs. PBS and of logFCs of ITDT vs. PBS, of genes upregulated in NK cells from a comparison of ITDT vs. PBS (up in ITDT), NS, or downregulated in a comparison of ITDT vs. PBS (down in ITDT). Significance threshold is based on adjusted p-value < 0.05 and abs(logFC) > 1. Top up and down genes based on Log FC are labeled on plot. (c) Percentages of NK cells (CD3-NKp46+) among splenocytes, splenocytes depleted of NK cells, or enriched NK cells. (d) Representative analysis of NK-depleted splenocytes and enriched NK cell populations. Cells were pregated for live+CD45+CD19-F4/80-Ter119- cells. (e) Cytotoxicity of MC38-*B2m^-/-^* target cells measured with 4-hour chromium release assays by splenocytes or NK cell-depleted splenocytes or enriched splenic NK cell populations as shown. Cells were prepared from MC38-*B2m^-/-^* tumor bearing mice treated as shown with DT (both IT and IP) 6 days previously, with or without IP depletion of CD4 T cells or with PBS alone. In the middle panel, splenocytes from these mice were tested after depleting NK cells in vitro, or not, before the assay. In the right panel, NK cells were enriched from all three samples. The target cell spontaneous release averaged 8.7% in these assays. (f) Baseline percentages of degranulation and IFNg production by NK cells without ex vivo stimulation with plate-bound NK1.1 antibodies. Data represent means ± SEM; *p < 0.05, **p < 0.01, ***p < 0.001, and ****p < 0.0001 from Ordinary one-way ANOVA with Dunnett’s multiple comparisons tests (a,c-d) or Welch’s t-tests (e-f). (n=5-8 mice/group, representative of at least 3 independent experiments).

**Ext. Data Figure 4.**
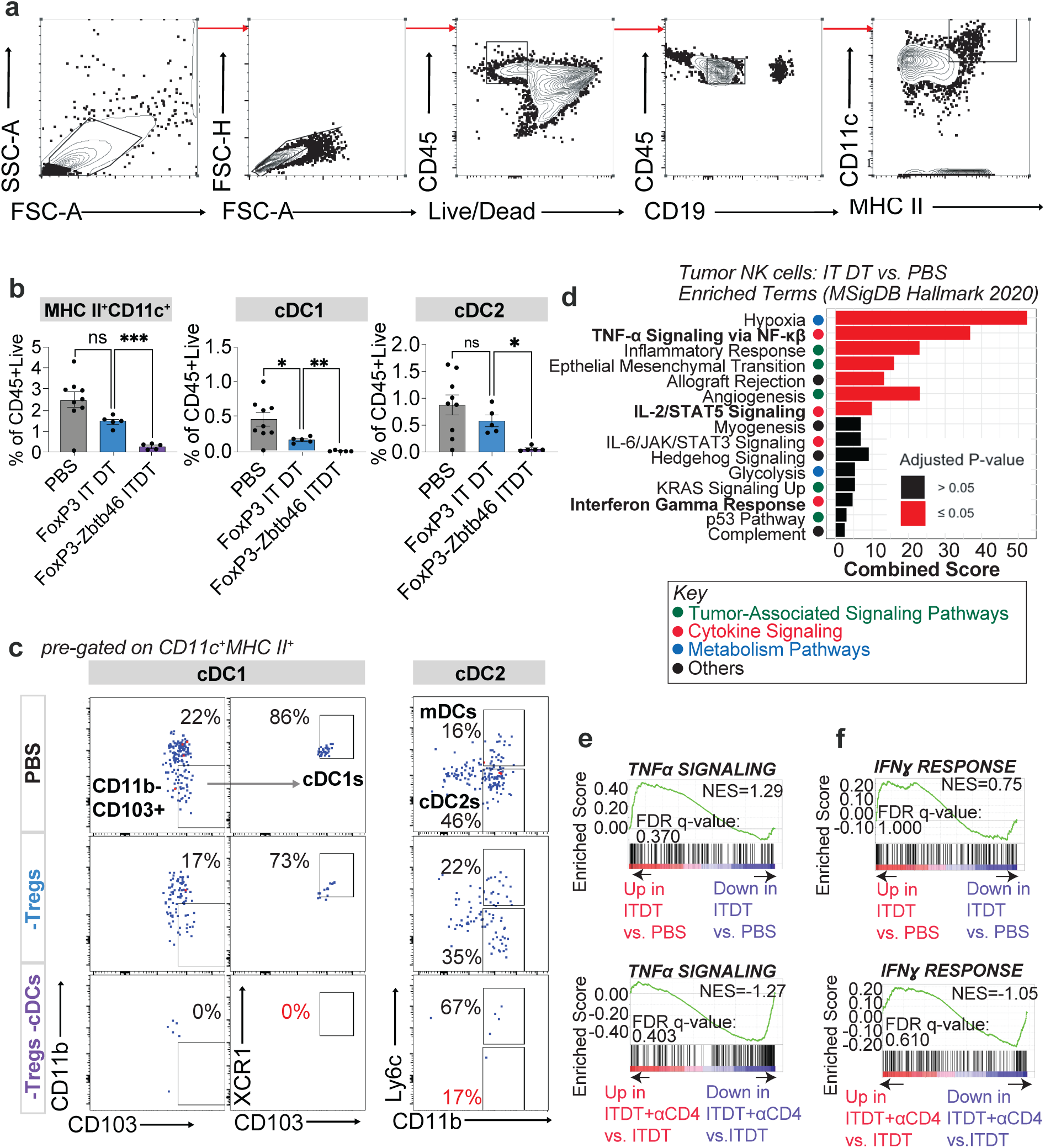
(associated with Fig 6): Bone marrow chimera validation and cytokine analyses (a) Representative flow gating strategy for bone marrow chimeras with *Foxp3^DTR^* and *Zbtb46^DTR^*;*Foxp3^DTR^* to characterize dendritic cells. (b) The bar graphs show CD11c+MHCII+ DCs as percentages of CD45+Live cells, DC1s (XCR1+CD103+CD11b-) and DC2s (CD11b+Ly6C-) in each treatment group with corresponding bar graphs pre-gated on Live+CD45+CD19-MHCII+CD11c+ as shown in S6A. One experiment performed. (c) Representative flow plots of DC1s and DC2s as shown in ext. data Fig 4b (d) Hypergeometric enrichment test, conducted with EnrichR with MSigDB Hallmark dataset. The top 15 pathways are shown and manually categorized as indicated by the circles and associated key. Combined score defined in EnrichR as the product of the odds ratio and -log(p), with p being the unadjusted p-value. (e-f)GSEA Enrichment plots for Hallmark TNF-alpha signaling and IFN-gamma signaling pathways upregulated in ITDT vs PBS NK cells or downregulated in ITDT+anti-CD4 versus ITDT NK cells FDR-q-value and normalized enrichment score (NES) are labeled in the plot. The Y-axes represent enrichment scores (ES). Data represent means ± SEM; *p < 0.05, **p < 0.01, ***p < 0.001, and ****p < 0.0001 from Ordinary one-way ANOVA with Dunnett’s multiple comparisons tests

**Ext. Data Figure 5.**
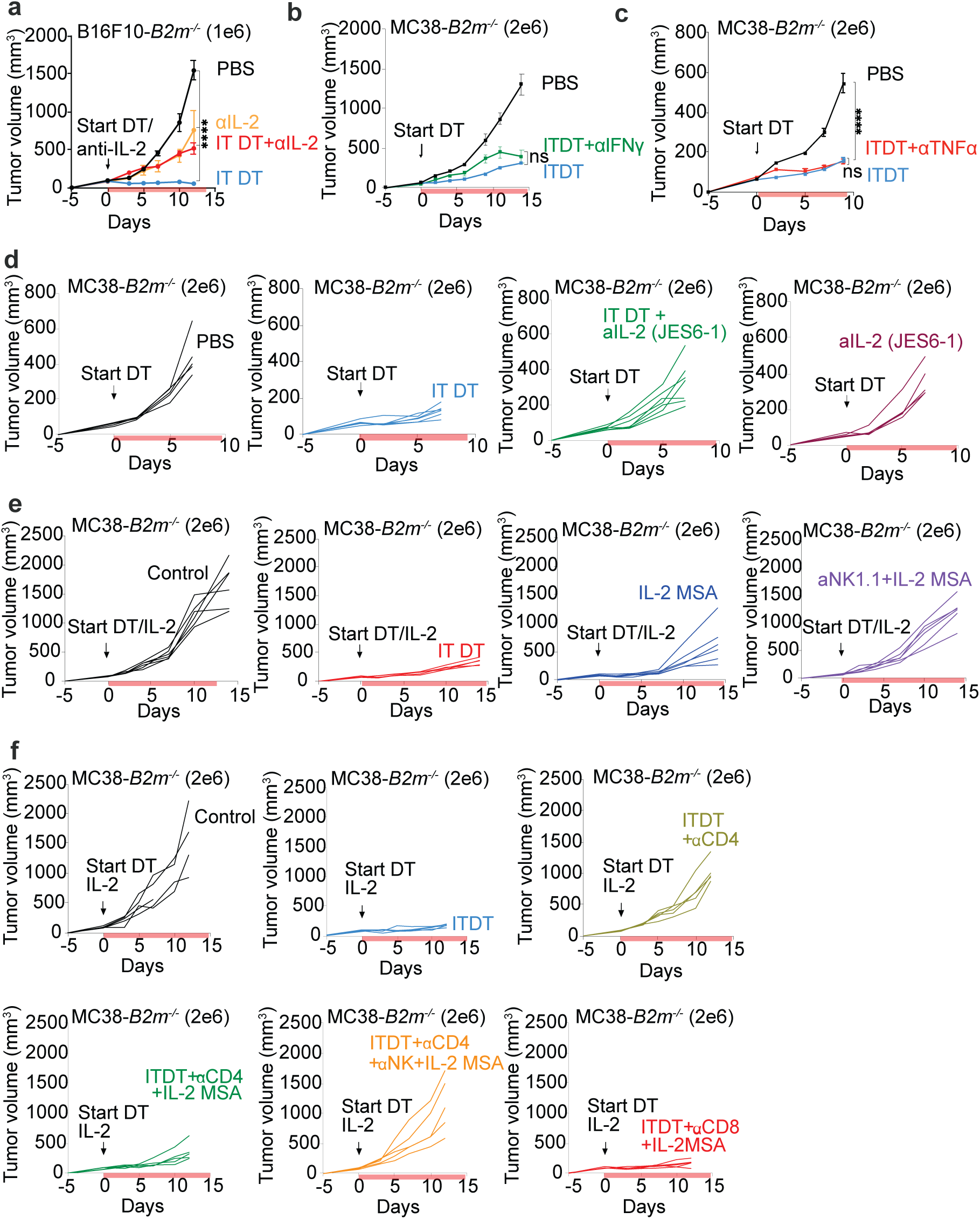
(associated with Fig 7): Tumor growth dependence on cytokines (a) Tumor growth curves of *Foxp3^DTR-GFP^* mice that were depleted of Tregs intratumorally with IT DT injections, or IT DT in combination with IP administration of anti-IL- 2 mAb JES6 (red), or with anti-IL-2 alone (yellow). One experiment was performed with B16F10*-B2m^-/-^* tumors. (b) Tumor growth curves of *Foxp3^DTR-GFP^* mice that were depleted of Tregs intratumorally with IT DT injections or in combination with anti-IFN*γ* (clone XMG1.2). MC38- *B2m^-/-^* tumors were tested. One representative experiment of two is shown. (c) Tumor growth curves of *Foxp3^DTR-GFP^* mice that were depleted of Tregs intratumorally with IT DT injections or in combination with anti-TNFα (clone 19.12). MC38-*B2m^-/-^* tumors were tested. One representative experiment of two is shown. (d) Tumor growth curves of *Foxp3^DTR-GFP^* mice that were depleted of Tregs intratumorally with IT DT injections. B16F10-*B2m^-/-^ Ifngr^-/-^* tumors were tested. One experiment was performed. (e-g) Spider plots of tumor growth shown in Figure 7. Data represent means ± SEM; *p < 0.05, **p < 0.01, ***p < 0.001, and ****p < 0.0001. RM two-way ANOVA followed by Geiser-Greenhouse correction

**Extended Data Figure 6:**
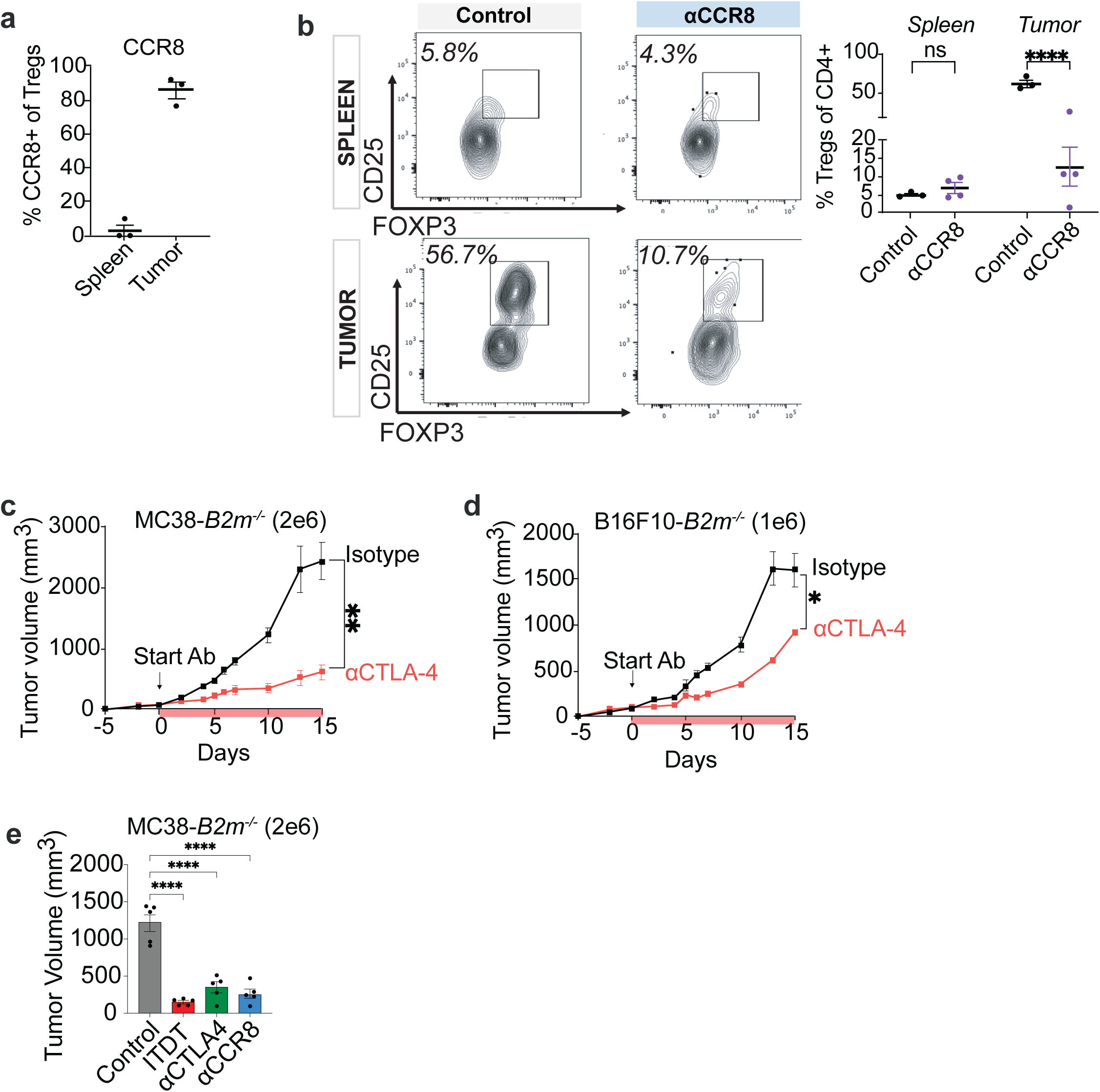
Alternatives to intra-tumoral Treg ablation ) Expression of CCR8 on Tregs in the spleen and tumor of MC38-*B2m^-/-^* tumor-bearing mice. One experiment performed. ) Representative flow plots and quantified frequencies of Tregs in spleen and tumor of mice treated with isotype control and anti-CCR8 (same experiment as in (A)-d) Growth curves of MC38-*B2m^-/-^* and B16F10-*B2m^-/-^* tumors in mice treated with isotype control IgG or CTLA-4 antibody. Representative experiments of 4 (MC38-*B2m*^-/-^) or 2 (B16F10-*B2m*^-/-^) performed are shown Data represent means ± SEM; *p < 0.05, **p < 0.01, ***p < 0.001, and ****p < 0.0001 Ordinary one-way ANOVA with Dunnett’s multiple comparisons tests. d: RM two-way ANOVA followed by Geiser-Greenhouse correction. (n=5-10 mice/group)

**Extended Data Figure 7:**
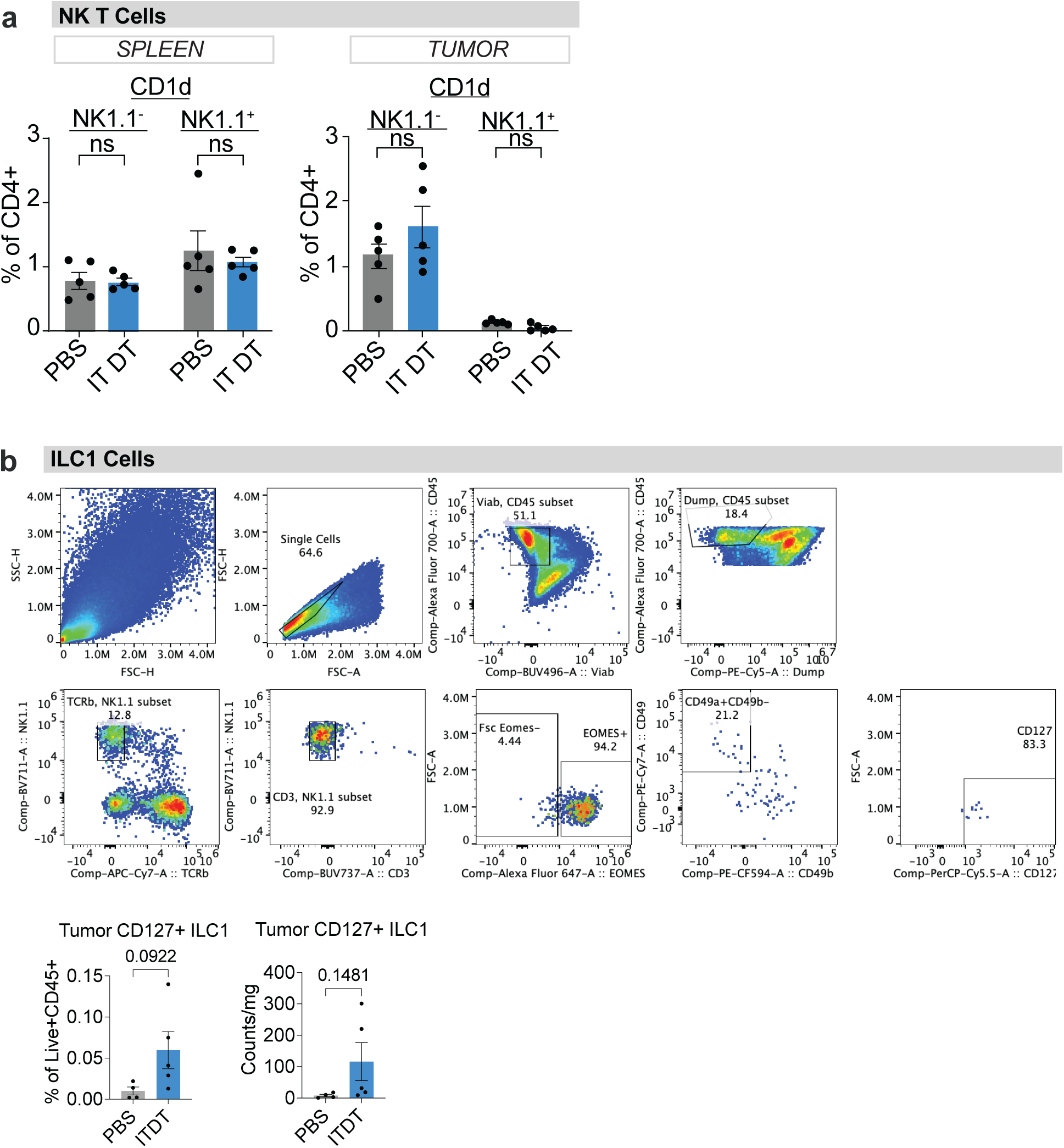
ILC1 cells and invariant NK1.1^+^ T cells in tumors and spleens (a)Percentages of invariant NKT cells detected by staining CD4 T cells with a- galactosylceramide loaded CD1d tetramers in spleens and tumors of IT DT treated mice compared to PBS-treated controls. Invariant NKT cells were rare in the tumors; note that even if they were more abundant, recognition of CD1d on these tumor cells cannot occur since functional CD1d expression is b2-microglobulin- dependent. (b-c) Gating strategy, percentages and counts of ILC1 cells in tumors of mice treated with either PBS or ITDT. Data represent means ± SEM; *p < 0.05, **p < 0.01, ***p < 0.001, and ****p < 0.0001 from Welch’s t-tests. (n=5-8 mice/group, one experiment)

